# A Reproducible Fetal Lamb Model of Complex Gastroschisis with Temporal Characterization of Bowel Changes

**DOI:** 10.64898/2026.03.25.714287

**Authors:** Tomohiro Arai, Michael A. Belfort, David Basurto, Marianna Scuglia, Kanokwaroon Watananirun, Wasinee Tianthong, Tom Bleeser, Marta Grinza, Simen Vergote, Emma Van den Eede, Michael Aertsen, Bethany Fisher, Alex Menys, Theo Thijs, Inge Depoortere, Alison Accarie, Ricard Farré, Tim Vanuytsel, Geert Molenberghs, Francesca Russo, Paolo De Coppi, Larry H. Hollier, Sundeep G. Keswani, Jan Deprest, Luc Joyeux

**Author notes:** **Corresponding authors Jan Deprest MD, PhD**, Department of Development and Regeneration, Biomedical Sciences, KU Leuven, Leuven, Belgium, Email address, **Luc Joyeux MD, PhD**, Texas Children’s Center for Translational Fetal Research, Texas Children’s Fetal Center, Texas Children’s Hospital, Houston, TX, USA. These authors contributed equally as first authors (Tomohiro Arai, Michael A. Belfort). These authors contributed equally as senior authors (Jan Deprest, Luc Joyeux).

## Abstract

**Objectives:** To determine whether a fetal lamb model reproducibly induces complex gastroschisis at term with its associated structural and functional bowel changes, and to compare its occurrence and associated bowel findings at different post-induction intervals.

**Methods:** Gastroschisis was surgically induced in fetal lambs at gestational day 75 by creating a standardized abdominal wall defect stabilized with a silicone ring. Fetuses were assessed at term or at mid-gestation (13–21 days after induction), using non-operated littermates as controls. The primary outcome was occurrence of complex gastroschisis (bowel stenosis, atresia, volvulus, perforation, or necrosis) in survivors. At mid-gestation, this was compared between early (13–16 days) and late (17–21 days) groups, using Fisher’s exact test. Sample size for the primary outcome at term was determined a priori (α=0.05, 80% power). Mid-gestation sample size was determined using the same statistical assumptions, informed by an initial exploratory cohort. Secondary outcomes included prenatal and in vivo imaging, ex vivo bowel contractility, amniotic fluid composition, and histology.

**Results:** Gastroschisis was induced in 32 fetuses. In the term cohort (n=14), 50% survived (7/14); all survivors had complex gastroschisis, with impaired bowel motility, altered enteric neural contractile responses, and smooth muscle remodeling compared with controls. Among fetuses assessed at mid-gestation (n=18), two died in utero; complex gastroschisis occurred more frequently in the late (5/7, 71%) than in the early (1/9, 11%; *p*=.04) group. Mid-gestation fetuses with gastroschisis had greater intra-abdominal bowel dilation on ultrasound and higher amniotic fluid digestive enzyme levels than controls, with the greatest dilation in fetuses with complex gastroschisis.

**Conclusions:** This fetal lamb model reproducibly induced complex gastroschisis with structural and functional bowel changes in surviving lambs at term. Complex gastroschisis occurred more frequently at 17–21 than at 13–16 days after induction. At mid-gestation, fetuses with gastroschisis showed greater bowel dilation and differences in amniotic fluid composition than controls, whereas structural and functional bowel changes were not detected.

## Introduction

Gastroschisis is a congenital abdominal wall defect in which the bowel eviscerates into the amniotic cavity. Approximately 20% of cases are classified as *complex* gastroschisis, characterized by bowel stenosis, atresia, volvulus, perforation, or necrosis at birth.^1–4^ Compared with simple gastroschisis, this phenotype is associated with substantially higher morbidity, including sepsis, bowel obstruction, and short bowel syndrome, as well as increased mortality.^3,4^

Ischemic and inflammatory mechanisms have been implicated in bowel injury in gastroschisis, potentially arising from mechanical constriction at the abdominal wall defect and bowel exposure to amniotic fluid, respectively.^5,6^ Clinical observations suggest that this process may be progressive during gestation.^7,8^ However, the sequence of bowel changes leading to the complex phenotype remains poorly defined.

Large-animal models enable assessment of disease development at defined gestational time points, including evaluation of bowel structure and function that are not feasible in humans. The fetal lamb is particularly well suited for translational investigation because of its anatomical similarity to the human fetus.^9^ However, prior experimental fetal lamb models of gastroschisis have reported variable phenotype generation, often in small cohorts without formal sample-size calculations.^10^ Understanding how complex gastroschisis develops requires a reproducible experimental model designed to consistently induce the defined anatomical features of complex gastroschisis, rather than reproduce its embryological origin or natural occurrence, while recapitulating its characteristic bowel features and the bowel dysfunction that is a major determinant of morbidity.^3,4^ Once such a phenotype is reliably established, studying bowel findings at different stages after induction may provide insight into how the phenotype develops within the experimental model.

Therefore, we aimed to determine whether this fetal lamb model reproducibly induces complex gastroschisis at term, together with its associated structural and functional bowel changes. We additionally aimed to compare the occurrence of complex gastroschisis and associated bowel findings at different stages after induction.

## Methods

Experiments were approved by the Ethics Committee on Animal Experimentation of KU Leuven (057/2021). All procedures complied with European Directive 2010/63/EU and Federation of European Laboratory Animal Science Associations guidelines. Reporting adhered to the Animal Research: Reporting of *In Vivo* Experiments 2.0 guidelines.^11,12^

### Study design

This study comprised two sequential phases. Gastroschisis was surgically induced at gestational day (GD) 75.^13,14^ In multiple pregnancies, only one fetus per litter underwent gastroschisis induction. In phase 1, fetuses were assessed at term (GD 135–140) to determine reproducibility of complex gastroschisis. In phase 2, fetuses were assessed 13–21 days after induction to evaluate temporal bowel changes.

The primary outcome was occurrence of *complex* gastroschisis, defined as bowel stenosis, atresia, volvulus, perforation, or necrosis at necropsy.^3,6^ In the absence of these findings, persistent bowel herniation was classified as simple gastroschisis. Intrauterine fetal deaths (IUFDs) were recorded, but gastroschisis type was not assigned. Non-operated littermates served as controls.

Secondary outcomes included prenatal ultrasound assessments (Doppler indices, intra-abdominal bowel dilation (IABD), extra-abdominal bowel volume (EABV)); in vivo magnetic resonance imaging (MRI)-based bowel function using the GIQuant motility score and gastrointestinal (GI) contrast score,^15,16^ and morphology (IABD, intra-abdominal bowel wall thickness (IABWT)); ex vivo bowel contractility and enteric neural responses; intestinal permeability (transepithelial electrical resistance (TEER) and fluorescein isothiocyanate-dextran 4 kDa (FITC-Dx4) flux); amniotic fluid composition (total protein, digestive enzymes, and L-lactate)^17–20^; and histology (epithelial and smooth muscle layer assessment). Birth weight was used to normalize histologic measurements.^21,22^ Fetal sex was not consistently recorded and was therefore not included as a biological variable. Detailed methods are provided in **Supplemental methods**, **Figure S1**, and **Figure S2**.

### Sample size

For phase 1, we hypothesized complex gastroschisis in at least 60% of gastroschisis-induced lambs and none in controls.^10,14^ Using a superiority design with a two-sided α=0.05 and 80% power, six surviving lambs per group were required.^23^

For phase 2, no prior data were available for sample size estimation. Therefore, an initial exploratory cohort of 10 fetuses informed expected complex gastroschisis rates of 20% (early, 13–16 days) and 80% (late, 17–21 days). Based on these assumptions, seven surviving fetuses per group were required (α=0.05, 80% power).^23^

### Surgical induction of gastroschisis

Time-dated ewes (term ≈145 days) underwent fetal surgery at GD 75 under general anesthesia (see **Supplemental methods**). Following maternal laparotomy and hysterotomy, a standardized 1-cm abdominal wall defect was created in the fetal lower abdomen. A 1 cm-diameter silicone ring was secured to the defect margins to maintain a standardized abdominal wall defect, and the bowel was exteriorized (**Figure 1A–B; Videoclip S1**). The fetus was returned to the uterus, intra-amniotic fluids and antibiotics were administered, and the uterus and maternal abdomen were closed in layers.

**Figure 1.**
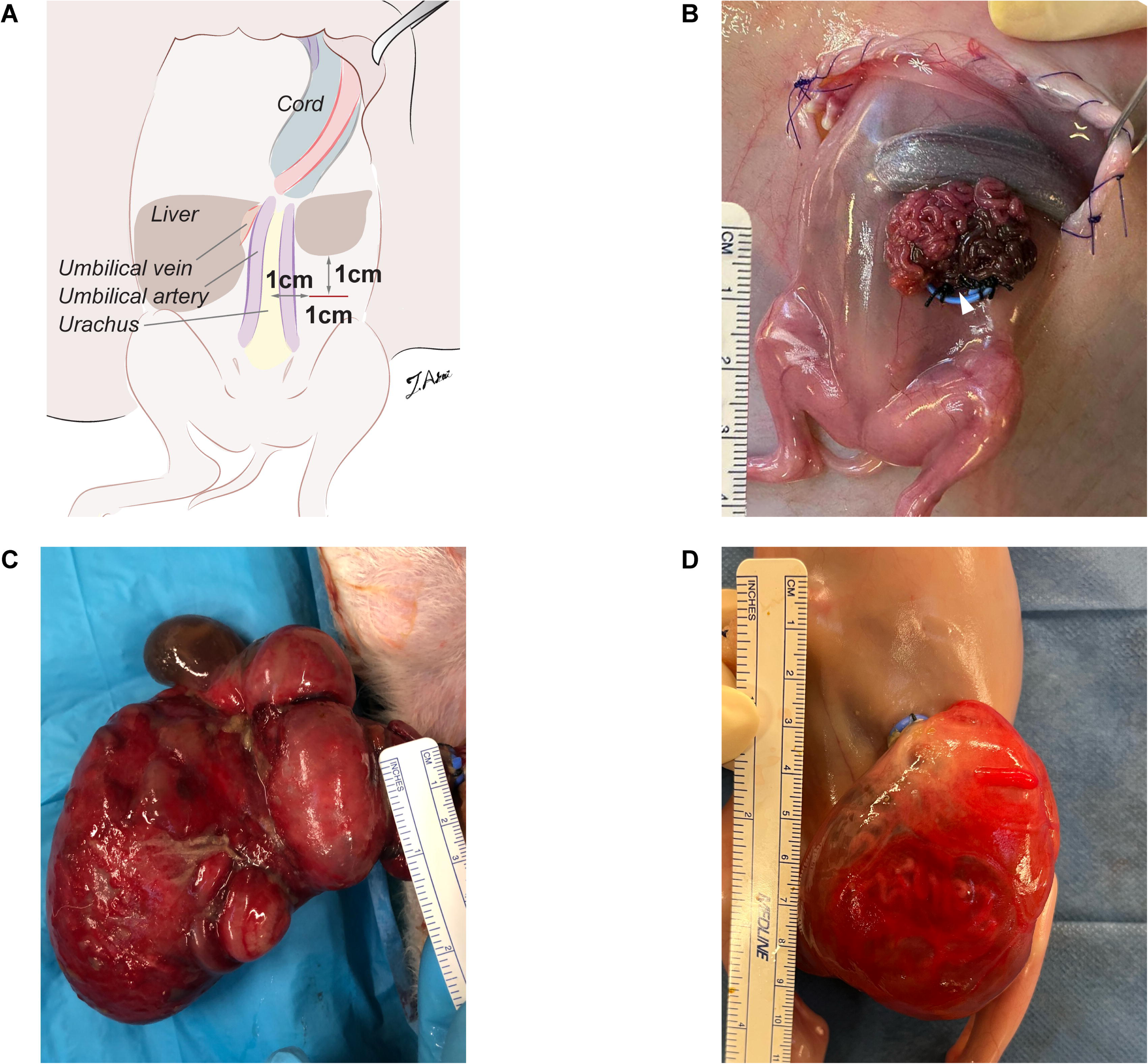
Surgical induction of gastroschisis in fetal lambs. At gestational day 75, following hysterotomy, the fetal lower abdomen and hindlimbs were exteriorized. (A) Schematic illustration of the surgical procedure. A 1-cm transverse full-thickness abdominal wall defect was created in the left lower quadrant, positioned 1 cm below the liver edge and 1 cm left of the midline. (B) Intraoperative photograph showing the silicone ring (blue) stabilizing the abdominal wall defect, with exteriorization of bowel to the proximal jejunum and distal spiral colon (arrowhead). (C) Representative term gastroschisis lamb demonstrating bowel dilation and a thick fibrotic peel surrounding the eviscerated bowel. (D) Representative mid-gestation gastroschisis fetus showing a thin translucent peel allowing visualization of the exteriorized bowel.

### Statistical analysis

GraphPad Prism (9.5.0 for macOS; GraphPad Software, La Jolla, CA, USA) was used for statistical analyses. The experimental unit was the individual fetal lamb. Analyses included only fetuses surviving to predefined assessment; IUFDs were recorded but excluded. The primary outcome was compared using two-sided Fisher’s exact tests (term: gastroschisis-induced lambs vs. non-operated littermate controls; mid-gestation: early vs. late).

The secondary outcomes were compared according to gastroschisis type at necropsy (i.e., complex gastroschisis, simple gastroschisis, or normal) either at term or at mid-gestation. Given small group sizes, normality was not assumed or formally tested. Data are presented as median (interquartile range) and compared using the Mann–Whitney U test for two-group comparisons or the Kruskal–Wallis test with Dunn’s post hoc multiple comparison test for three-group comparisons. For secondary-outcome biomarker analyses of prenatal ultrasound and amniotic fluid parameters, comparisons were performed first between pooled gastroschisis and normal groups, then between complex and simple gastroschisis, using the Mann–Whitney U test. As an exploratory analysis, logistic regression with receiver operating characteristic (ROC) curve analysis was used to determine an optimal IABD cut-off value for predicting complex gastroschisis, based on Youden’s index. A two-sided *p*<.05 was considered statistically significant.

## Results

Gastroschisis was induced at median GD 75 in 32 fetuses (see **Figure 2** for experimental timeline and group allocation).

**Figure 2.**
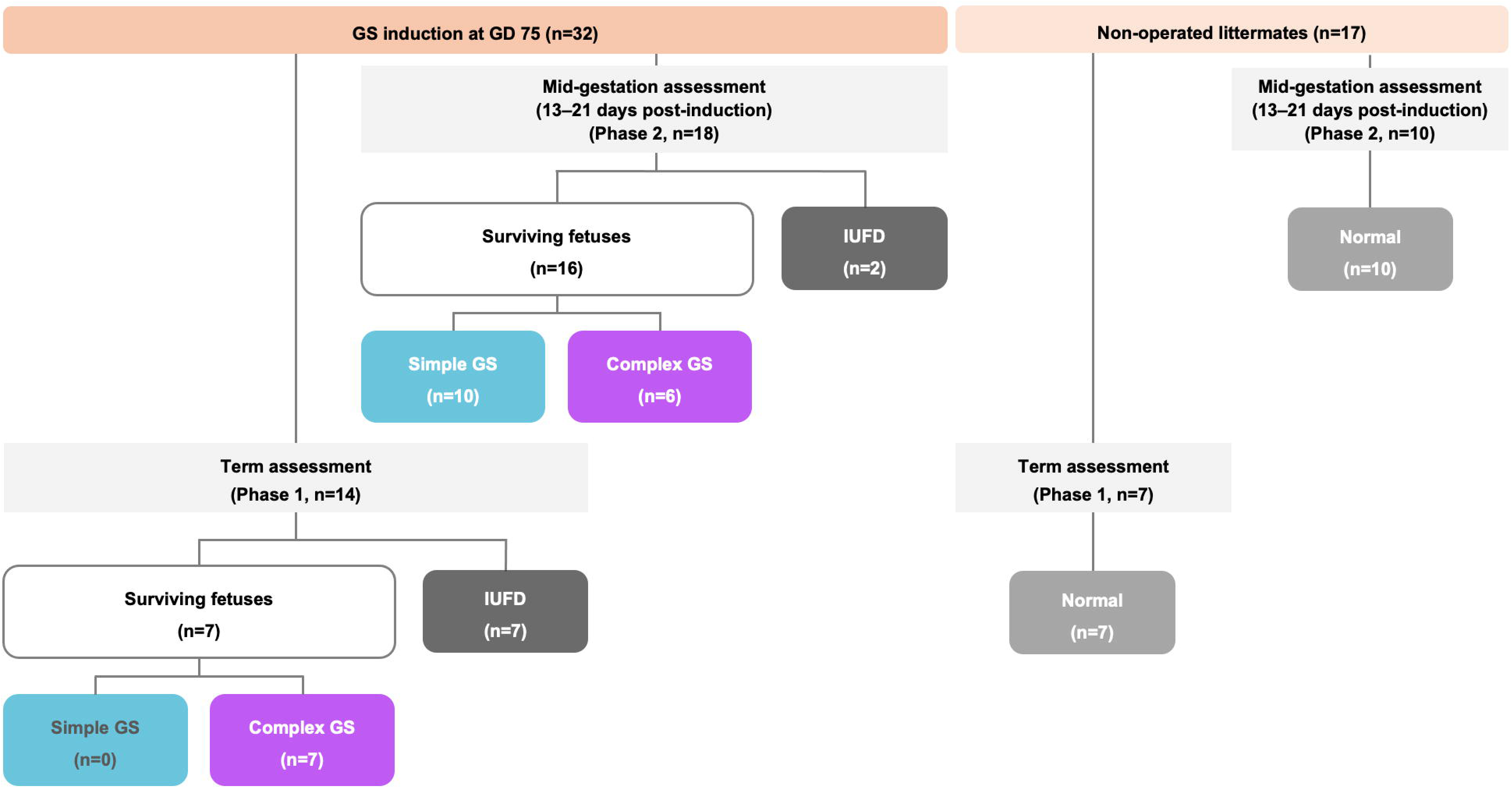
Experimental flowchart. Gastroschisis was surgically induced in fetal lambs at gestational day 75. Fetal lambs were assessed either at term (phase 1) or at 13–21 days post-induction (phase 2). Non-operated littermates served as controls at corresponding time points. Abbreviations: GD, gestational day; GS, gastroschisis; IUFD, intrauterine fetal death.

### Phase 1: Reproducibility of complex gastroschisis

At term (63–69 days post-induction; GD 138–144), survival after induction was 50% (7/14). All surviving lambs (7/7) exhibited complex gastroschisis (**Figure 1C**; **Table 1**). All non-operated littermates (7/7) survived and had normal-appearing bowel. Complex gastroschisis was most frequently characterized by bowel stenosis at the level of the silicone ring (6/7; **Figure S3A**), with atresia, perforation, and necrosis each observed in one lamb. Volvulus was not observed. All IUFD fetuses were macerated at assessment, although one case had findings suggestive of volvulus (**Figure S3B**; Table S1).

**Table 1.** Primary outcome: Gross anatomical phenotype at assessment after induction.

| Group | Complex | Simple | IUFD | Normal | Total |
| --- | --- | --- | --- | --- | --- |
| Term lambs |  |  |  |  |  |
| Gastroschisis-induced | 7 | 0 | 7 | 0 | 14 |
| Non-operated littermates | 0 | 0 | 0 | 7 | 7 |
| Mid-gestation fetuses (13–21 days post-induction) |  |  |  |  |  |
| Gastroschisis-induced, 13–16 days | 1 | 8 | 1 | 0 | 10 |
| Gastroschisis-induced, 17–21 days | 5 | 2 | 1 | 0 | 8 |
| Non-operated littermates | 0 | 0 | 0 | 10 | 10 |

Values represent numbers of fetuses. Two-tailed Fisher's exact test showed a significant difference in complex gastroschisis occurrence among survivors at term between gastroschisis-induced and non-operated littermates ( $p<.001$ ), and between early (13–16 days) and late (17–21 days) post-induction groups ( $p=.04$ ).
Abbreviations: IUFD, intrauterine fetal death.

The availability of secondary outcome measurements across groups is summarized in Table S2A. On in vivo MRI, gastroschisis-induced lambs exhibited lower GIQuant motility scores, higher GI contrast scores, greater IABD, and increased IABWT compared with controls (*p*=.004, *p*<.001, *p*=.004, and *p*=.004, respectively; **Figure 3A–D**). Ex vivo bowel smooth muscle contractile responses to acetylcholine, potassium chloride, and substance P were not significantly different between groups. Neural contractile responses to electrical field stimulation (EFS) required higher stimulation frequencies to reach half-maximal responses in gastroschisis-induced lambs (*p*=.02; **Figure 3E**), whereas maximal contractions were comparable between groups. Pharmacological analyses demonstrated no difference in overall non-adrenergic, non-cholinergic responses, a reduced L-NAME-associated effect (*p*=.001), and no difference in the MRS-2500-associated effect compared with controls (**Figure S4A**). In ex vivo intestinal permeability testing, both TEER and FITC-Dx4 flux changed over time but did not significantly differ between groups (**Figure S5**). Histologic analysis of the terminal ileum demonstrated increased muscular thickness (*p*=.005; **Figure 3F**) and reduced muscular cell density in gastroschisis-induced lambs, with similar findings across proximal, middle, and distal bowel segments (**Figure S6**).

**Figure 3.**
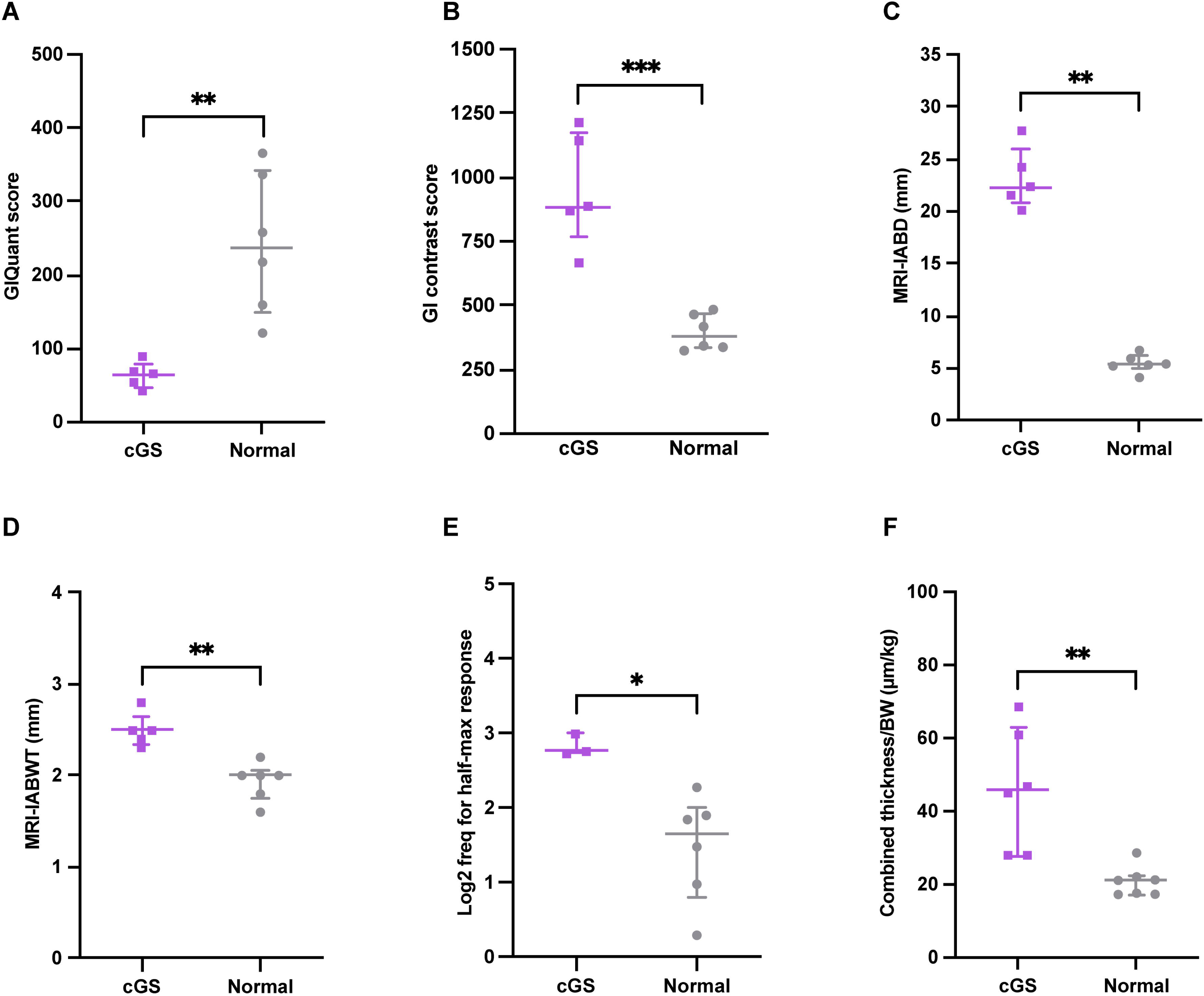
Functional and structural bowel alterations in term lambs. (A–D) In vivo MRI parameters: (A) GIQuant motility score, (B) GI contrast score, (C) IABD, and (D) IABWT. (E) Ex vivo bowel contractility testing: log_2_-transformed stimulation frequency inducing half-maximal neural response. (F) Histologic bowel morphology of the terminal ileum: body-weight-adjusted combined muscular-layer thickness (circular + longitudinal). \**p*<.05; \*\**p*<.01 Abbreviations: BW, birth weight; cGS, complex gastroschisis; freq, frequency; IABD, intra-abdominal bowel dilation; IABWT, intra-abdominal bowel wall thickness; MRI, magnetic resonance imaging.

### Phase 2: Duration required for complex gastroschisis development after induction

Of 18 induced fetuses assessed at mid-gestation (**Figure 1D**), ten were assessed at 13–16 days and eight at 17–21 days post-induction. One IUFD occurred in each group, leaving nine and seven surviving fetuses, respectively. All non-operated littermates (10/10) survived and had normal-appearing bowel. Among survivors, complex gastroschisis occurred in one lamb in the early group (1/9) and five lambs in the late group (5/7), resulting in a higher occurrence in the late group than in the early group (*p*=.04; Table 1). Among the six complex gastroschisis cases assessed at mid-gestation, atresia (5/6) and perforation (3/6) predominated (**Figure S3C–D**), whereas stenosis was observed in one fetus at 13 days after induction. In IUFD cases, further characterization of eviscerated bowel was not possible due to maceration (Table S1).

The availability of secondary outcome measurements across groups is summarized in Table S2B. Ex vivo bowel smooth muscle contractility and neural contractile responses, with or without pharmacological blockers, did not differ significantly among groups (**Figure S4B**). Histologic analyses demonstrated no differences among groups across proximal, middle, and distal bowel segments (**Figure S7**).

On prenatal ultrasound (**Figure S8**), when all surviving gastroschisis fetuses were compared with controls, only IABD differed (*p*=.009). Within gastroschisis fetuses, IABD was greater in complex than in simple gastroschisis (*p*=.04). ROC analysis indicated an IABD threshold of 2.8 mm for association with complex gastroschisis among gastroschisis-induced fetuses (area under the curve, 0.925; 95% CI, 0.743–1.000; **Figure S9**). EABV did not differ between complex and simple gastroschisis, although all perforation cases exhibited EABV<10 cm^3^. In amniotic fluid, total protein, lipase, and L-lactate were higher in gastroschisis than in controls (*p*=.009, *p*=.03, and *p*=.04, respectively; **Figure S10**). L-lactate was lower in complex than simple gastroschisis (*p*=.03).

A summary of the principal findings across the studied gestational time points is provided in **Figure 4**.

**Figure 4.**
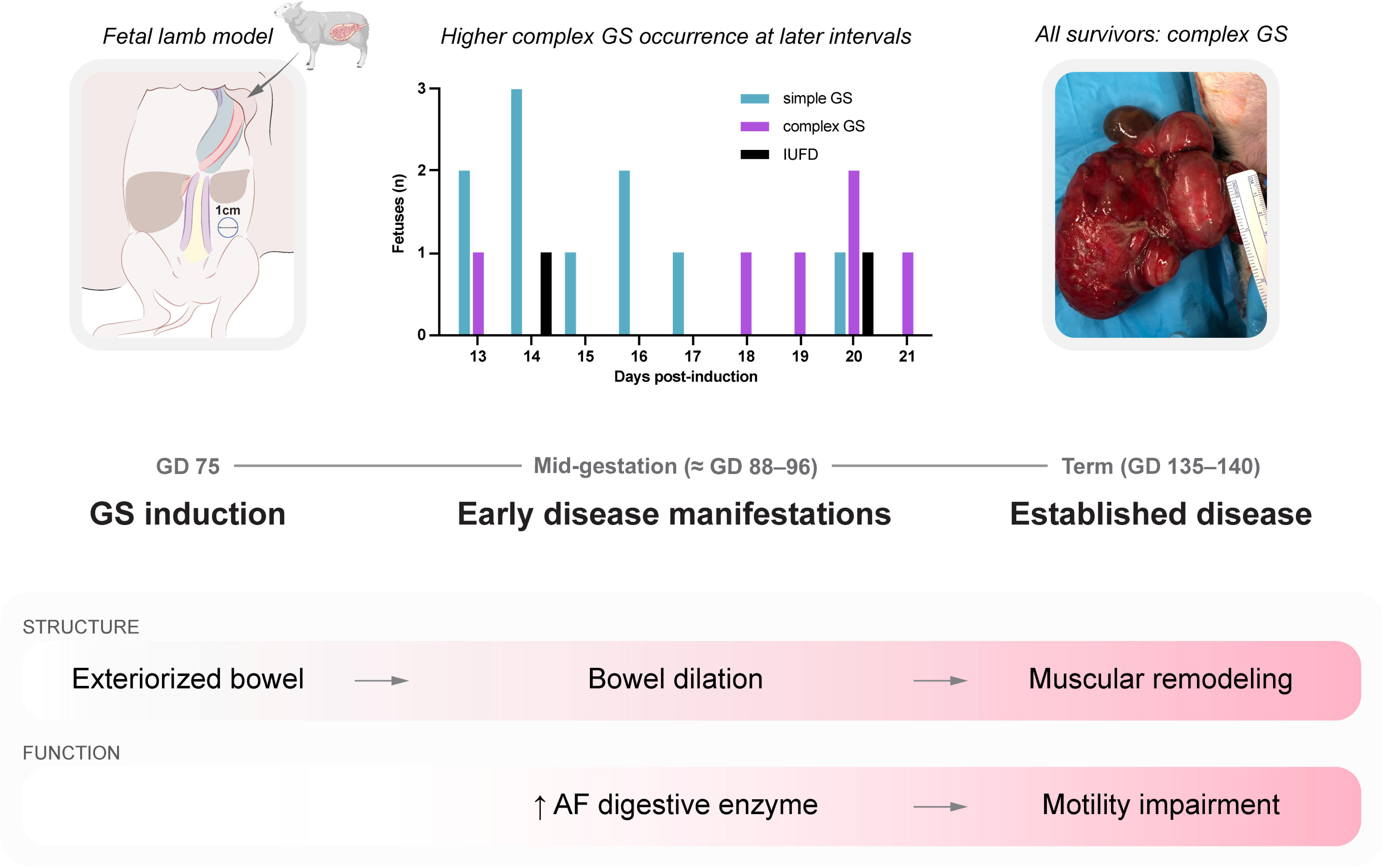
Proposed temporal development of complex gastroschisis in the fetal lamb model. Gastroschisis was induced in fetal lambs at gestational day 75. During mid-gestation, complex gastroschisis occurred more frequently at later post-induction intervals. Early changes were characterized by bowel dilation and increased amniotic fluid digestive enzyme levels, suggesting impaired intestinal transit. These alterations were followed, in independently assessed term cohorts, by smooth-muscle remodeling and motility impairment, as well as consistent occurrence of anatomical features of complex gastroschisis at term. Sheep illustration created with BioRender.com. Abbreviations: AF, amniotic fluid; GD, gestational day; GS, gastroschisis; IUFD, intrauterine fetal death.

## Discussion

We describe a fetal lamb model with reproducible complex gastroschisis. At term, all surviving lambs developed complex gastroschisis, with structural changes consistent with smooth muscle remodeling and functional changes in bowel transit, motility, and enteric neural responses. At mid-gestation, complex gastroschisis occurred more frequently at 17–21 than at 13–16 days after induction. At that stage, bowel dilation and changes in amniotic fluid composition were present, whereas the structural and functional changes, observed at term, were not.

The silicone ring used to stabilize the abdominal wall defect was a deliberate choice. Previous ovine gastroschisis models frequently exhibited spontaneous bowel reduction and defect closure, or inconsistent defect size at term,^10^ limiting reproducible generation of complex gastroschisis features. Notably, complex gastroschisis was only observed when a stabilizing silicone ring was used.^14^ We therefore deliberately chose this model, as we only aimed to study complex gastroschisis. This was associated with an IUFD rate of 50%, substantially higher than the ∼4.5% reported for all forms of gastroschisis.^24^ Individual causes of IUFDs could not be determined because of lack of longitudinal monitoring and maceration, although body weight at necropsy suggested some deaths occurred far beyond induction of the defect. High IUFD rates have been reported in earlier experiments.^14,25^ This may be a limitation of the model; nonetheless, the complex gastroschisis rate among survivors met the study objective.

Of great interest is that our model also recapitulates structural and functional bowel changes typical for complex gastroschisis. Structurally, complex gastroschisis lambs had increased bowel wall thickness on in vivo MRI, which was corroborated on histology by increased muscular-layer thickness, consistent with previous ovine models.^17,19,26^ Although the terminal ileum is considered most susceptible to injury,^21,22,27^ similar changes occurred across all bowel segments in the present model. Functionally, enteric neural responses were altered, with higher stimulation frequencies required to reach half-maximal contractile responses on EFS, aligning with impaired contractility reported previously.^19,25^ However, responses to acetylcholine, potassium chloride, and substance P did not differ between groups in contrast to earlier studies,^19,25^ although none of these studies was powered for this secondary outcome. Conversely, bowel transit was impaired, evidenced by higher GI contrast scores, corroborating previous postnatal observations in complex gastroschisis lambs.^14^ Motility was further reduced on GIQuant analysis, a previously described quantitative MRI-based motility measure.^15^ Given the clinical importance of bowel dysfunction in complex gastroschisis,^3,4^ these functional changes support the model’s translational value.

At mid-gestation, atresia and perforation predominated, whereas stenosis, the dominant lesion at term, was uncommon. Whether these lesions represent different stages of a common pathological process, heterogeneous pathways to different presentations, or coincidental findings cannot be determined from our data. Structural remodeling and bowel dysfunction observed at term were not detected at mid-gestation. Contractility findings should be interpreted cautiously given limited numbers, whereas the larger histologic sample size suggests that structural remodeling was not yet detectable. Of interest is that ultrasound and amniotic fluid biomarkers already discriminated affected fetuses. In humans, increased second-trimester IABD has been associated with complex gastroschisis at birth.^7,28,29^ In our model, increased IABD coincided with macroscopic complex gastroschisis at mid-gestation, suggesting it may reflect concurrent disease status rather than solely predict later disease. Total protein, lipase, and L-lactate were elevated in gastroschisis as well, consistent with previous fetal lamb studies.^17–19^ L-lactate is not bowel-specific and may reflect systemic fetal metabolism rather than localized bowel injury.^30^ Unexpectedly, L-lactate was higher in simple than in complex gastroschisis, which remains unexplained.

Overall, our mid-gestation findings suggest that prenatal imaging and biochemical changes are detectable before the full-blown structural and functional phenotype observed at term. Although our study was not mechanistic in nature, the structural and functional changes observed can be explained through several mutually non-exclusive mechanisms, including ischemia and inflammation.^5,6^ Molecular analysis of a partly overlapping cohort identified extracellular matrix remodeling, angiogenesis, inflammation, and enteric nervous system changes, providing complementary and hypothesis-generating mechanistic context.^31^ Future models designed to distinguish ischemic from inflammatory injury will be required to clarify their relative contributions.^19^

Across both gestational time points, our data suggest that anatomical features of complex gastroschisis are present before the structural and functional changes observed at term (**Figure 4**). This forms the basis of our current integrated view of complex gastroschisis as a process comprising distinct anatomical, structural, and functional components. This model therefore provides a platform for future mechanistic and interventional studies to better understand, and ultimately modify, disease progression.

Strengths of this study include the reproducible nature of the condition in a large-animal model, identification of a 2–3-week post-induction interval for further studying disease progression and potential fetal interventions, and adequate power for the primary outcome.^10^ Bowel structure and function were comprehensively assessed using complementary imaging, functional, biochemical, and histologic approaches. A key limitation is the thick fibrotic peel covering the exteriorized bowel from 13 days onward and universally present at term. This differs from the typical clinical presentation of free-floating bowel with localized peel formation,^32^ yet similar bowel matting may rarely occur.^33^ This feature is consistently reported in ovine studies^13,19,34^ and attributed to the high hyaluronic acid content of ovine amniotic fluid,^35^ but its biological impact remains unclear and may limit direct translatability. Because no surviving animals exhibited simple gastroschisis at term, whether the structural and functional changes are specific to complex gastroschisis or reflect gastroschisis more generally cannot be determined. Consequently, this model cannot address questions specific to *simple* gastroschisis. Some outcomes were not available because of IUFD, insufficient sample volume, apparatus availability, or logistical constraints. Although secondary outcomes were prespecified, they were not individually powered and should therefore be interpreted cautiously. Finally, the mid-gestation sample size was informed by an initial exploratory cohort, and the 2.8-mm IABD cutoff was also exploratory.

## Conclusions

This study describes a reproducible large-animal model in which all surviving lambs showed complex gastroschisis at term. Complex gastroschisis occurred more frequently at 17–21 than at 13–16 days after induction. Bowel dilation and changes in amniotic fluid composition were present at mid-gestation, whereas structural and functional bowel changes were detected at term.

## Acknowledgments

This work was supported by internal funding from Texas Children’s Hospital (LJ), the Men of Distinction Award (SK, MB), the Karakin Foundation (SK, MB), and the Texas Children’s Auxiliary Award (LJ). Jan Deprest is supported by the Great Ormond Street Hospital for Sick Children’s Charity. Tomohiro Arai is supported by the Funai Overseas Scholarship. Paolo De Coppi is supported by National Institute for Health Research (NIHR-RP-2014-04-046), Great Ormond Street Hospital Children’s Charity, and the NIHR Biomedical Research Centre at Great Ormond Street Hospital for Children NHS Foundation Trust. The funding sources had no role in study design, data collection, analysis, interpretation, or manuscript preparation. The authors declare that they have no competing interests.

We thank Sander Van den Broucke from the Center for Surgical Technologies, KU Leuven, Belgium, for assistance with animal handling, and Katrien Luyten from the G-PURE laboratory for assistance with histologic processing. We are grateful to Glynis Frans (Department of Laboratory Medicine, University Hospitals Leuven) and Sofie Vanmechelen for technical support and guidance in the biochemical analysis of amniotic fluid samples.

## Generative AI disclosure

During manuscript preparation, the authors used ChatGPT (OpenAI, 1.2026.184, OpenAI, San Francisco, CA, USA) and Claude (Anthropic, 1.52386.6, Anthropic PBC, San Francisco, CA, USA) for language polishing and light editorial assistance. The authors reviewed and approved all changes and take full responsibility for the manuscript content.

