## supplements for "A Reproducible Fetal Lamb Model of Complex Gastroschisis with Temporal Characterization of Bowel Changes"

**Supplementary methods**

**Animals and anesthesia**
Time-dated ewes (Swifter, term = 145 days) were provided by TRANSfarm, the large-animal facility of KU Leuven. The ewes were transferred at least 7 days before the first intervention to acclimate to the experimental facility. Upon arrival, pregnancy and fetal cardiac activity were confirmed by ultrasound before enrolment into the study. They were randomly housed, and caregivers were blinded to group allocation and the experimental protocol. Ewes were fasted for food, but not water, for 12 hours before surgery.
Anesthesia was induced in a transport cart by inhalation of 8 Vol-% sevoflurane (Sevotek, Fendigo, Brussels, Belgium) in 100% oxygen via a face mask. After loss of consciousness, sevoflurane was decreased to 4 Vol-%, and the ewe was transferred to the operating table and placed in a left lateral position. A pulse oximeter was attached to its ear, and an 18-gauge catheter (BD Insyte-W, BD, Sandy, UT, USA) was inserted into the right jugular vein. Following intravenous buprenorphine (0.02 mg/kg; Vetergesic Multidose, Ecuphar, Oostkamp, Belgium) and propofol (3 mg/kg; PropoVet Multidose, Ecuphar), endotracheal intubation was performed using direct laryngoscopy with a 9-mm internal diameter tube. Mechanical ventilation was provided using an Aespire S/5 anesthesia engine (Datex-Ohmeda, Madison, WI, USA) with 30–35% inspiratory oxygen, an initial tidal volume of 8 mL/kg, and a respiratory rate of 20/min, titrated to achieve an arterial pCO_2_ of 30 mmHg and a positive end-expiratory pressure of 5 cmH_2_O. Sevoflurane was maintained at an end-tidal concentration of 5 Vol-%. Correct endotracheal tube position was confirmed by bilateral vesicular breathing sounds, thorax excursions, tube condensation, and capnography. A 22-gauge catheter (BD Insyte-W, BD, Sandy, UT, USA) was placed in the ear artery for invasive blood pressure monitoring. Standard American Society of Anesthesiologists monitoring was applied (Datex-Ohmeda S/5 monitor, Datex-Ohmeda, Helsinki, Finland), including electrocardiography, invasive arterial blood pressure, pulse oximetry, end-tidal CO_2_, and inspiratory and expiratory oxygen and sevoflurane concentrations. Fetal heart rate was measured using ultrasound before and after surgery. A 1.27-cm-diameter gastric tube and an esophageal temperature probe were placed. Maternal body temperature was maintained between 38.4 and 40.0°C using a forced-air warming system (Bair Hugger, Arizant Healthcare, Eden Prairie, MN, USA).^1^ Mean arterial pressure was maintained above 67 mmHg (80% of awake value^2^) using intermittent peripheral intravenous infusions of noradrenaline (200 µg/mL; Noradrenaline, Aguettant, Lyon, France). At skin incision, 2.0 g cefazolin (CeFAZolin Sandoz, Sandoz Novartis, Holzkirchen, Germany) and a second dose of buprenorphine (0.02 mg/kg) were administered. During laparotomy, Hartmann solution was infused at 5 mL/kg/hour. Arterial blood gas analysis was performed to adjust ventilation, followed by lung recruitment maneuvers (40 cmH_2_O) at 10 and 75 min after intubation. During skin closure, sevoflurane administration was discontinued halfway through the closure, and 100% oxygen was administered. Postoperatively, 150 mg medroxyprogesterone acetate (Depo-Provera, Pfizer, Brussels, Belgium) was administered subcutaneously as a tocolytic, and 20 mL of 0.25% levobupivacaine (Levobupivacaine, Fresenius Kabi, Utrecht, Netherlands) was infiltrated subcutaneously into the incision for local analgesia. All monitoring equipment, except the pulse oximeter and intravascular lines, were removed, and the ewe was extubated once spontaneous breathing and purposeful movements were observed.
After recovery, ewes were returned to their stable with free access to food and water and were monitored daily for pain and wellbeing. Once the ewe began standing and feeding, buprenorphine (0.02 mg/kg) was administered intramuscularly. Additional postoperative analgesia and tocolysis were provided with intramuscular meloxicam (0.5 mg/kg; Metacam, Boehringer Ingelheim, Ingelheim am Rhein, Germany) at 24 hours postoperatively.
Postoperative follow-up consisted of daily maternal monitoring. In the event of maternal abnormalities, including reduced appetite or decreased physical activity, a standardized clinical assessment was performed, comprising measurement of body temperature, auscultation, and ultrasound confirmation of fetal viability. In cases of intra-uterine fetal death (IUFD), the fetus was collected according to the study protocol and necropsy was performed.

**Surgical induction of gastroschisis**

Before induction surgery, fetal cardiac activity was confirmed by ultrasound. Time-dated ewes (term ≈ GD 143–145) underwent surgery at GD 75 under general anesthesia. After maternal midline laparotomy (13 cm), a 6 cm hysterotomy was performed, and the membranes were secured with a purse-string suture using 3-0 polydioxanone (PDS II, Ethicon, Zaventem, Belgium). The fetus received intramuscular fentanyl (0.01 mg/kg; Fentadon, Dechra, Bladel, the Netherlands), and the lower abdomen and hindlimbs were exteriorized.
A standardized abdominal wall defect was created in the left lower quadrant using a 1-cm transverse laparotomy positioned 1 cm below the liver edge and 1 cm lateral to the midline to avoid liver and umbilical artery injury. A 1-cm diameter silicone ring was secured to the defect margins using four interrupted 2-0 silk sutures (Mersilk, Ethicon) to stabilize the defect. The small and large bowel were gently exteriorized using cotton-tipped applicators until both the proximal jejunum and spiral colon were located outside the abdominal cavity.
The fetus was returned to the uterus, after which 200 mL Ringer’s lactate and 500 mg cefazolin (Cefazolin Sandoz, Sandoz Novartis, Holzkirchen, Germany) were instilled into the amniotic cavity. The hysterotomy was closed with a running 3-0 polydioxanone suture, followed by closure of the maternal abdomen in two layers using size 1 polydioxanone for the fascia and 2-0 poliglecaprone (Monocryl, Ethicon) for the skin.

**Prenatal ultrasound**
Ultrasound measurements were done in all mid-gestation fetal lambs using a Voluson E10 system (GE Healthcare, Zipf, Austria) equipped with an RM6C transducer (2.0–6.0 MHz). Scanning was performed under general anesthesia through the exteriorized uterus while maintaining placental circulation, immediately prior to fetal collection.
The following parameters were recorded: pulsatility index (PI) of (a) umbilical artery (UA), (b) ductus venosus (DV), and (c) cranial mesenteric artery (CMA; corresponding to the superior mesenteric artery in humans); (d) width of bowel dilation outside the abdomen, i.e. extra-abdominal bowel dilatation (EABD), (e) width of bowel dilation inside the abdomen, i.e. intra-abdominal bowel dilation (IABD), (f) extra-abdominal bowel wall thickness (EABWT), (g) intra-abdominal bowel wall thickness (IABWT), and (h) extra-abdominal bowel volume (EABV).19-22 For the Doppler acquisitions, settings were as follows: magnification, sample volume 0.5–1.0 mm, within 30 degrees angle, and wall motion filter 50–70.**^3^**
All measurements were performed offline using the Voluson E10 BT16 software (EC310, GE Healthcare) on recorded videos and images to minimize anesthesia time. Parameters were measured three times and averaged. For the bowel measurements (i.e., EABD, IABD, EABWT, IABWT, and EABV), the bowel was distinguished from other duct-like structures such as vessels, urinary bladder, bile ducts, and gallbladder with frame-by-frame playback of the recordings. Then, gain of the B-mode was decreased as much as possible. Bowel dilation was defined as the distance between the inner edges of the bowel wall (i.e., lumen). For the EABD, measurements were taken in the transverse plane in the three most dilated bowel segments. Measurements for the IABD were made in the three most dilated bowel segments near the stalk of the exteriorized bowel, also in the transverse plane. Bowel wall thickness was defined as the distance between the outer and inner edges of the bowel wall. Measurements were obtained using the same images for EABD and IABD to maintain consistency in anatomical planes and reference points. Inter- and intra-rater reliability for the bowel measurements was assessed prior to analysis and is reported below. For the EABV estimation, three orthogonal diameters of the external bowel peel were measured. Length and width were defined in the transverse plane, while height was defined as the maximal cranio-caudal diameter measured perpendicular to the transverse plane. Volume was calculated using the ellipsoid formula: EABV = 1/2 (i.e., approximation of π/6) x length x width x height.

**Ultrasound measurement reliability (quality control)**
Inter- and intra-rater reliability was assessed for IABD, IABWT, EABD, and EABWT prior to analysis. Two raters independently measured a predefined set of stored images, and one rater repeated measurements on the same images. Reliability was summarized using correlation estimates (ρ) with 95% confidence intervals calculated by Fisher’s z transform (SAS 9.4; SAS Institute, Cary, NC, USA). Study measurements were performed according to the standardized measurement protocol described above. Reliability results are summarized in **Table S3**.

**Delivery and euthanasia**
*Term* lambs were delivered by cesarean section under loco-regional anesthesia. Before commencement of the terminal procedure, fetal cardiac activity was confirmed by ultrasound. The ewes were shortly sedated by mask inhalation of 8 Vol-% sevoflurane. Once required level of sedation was achieved, it was decreased to 2 Vol-%. In the left lateral position, an 18-gauge catheter was inserted into the right jugular vein, and a blood sample was drawn. Then in the prone position, spinal anesthesia was performed at the level of L6–S1 with 0.15 mL/kg lidocaine 2% (Lignocaine HCl 2% + Adrenaline tartr. 0.0036% KELA, Kela, Hoogstraten, Belgium).^4^ Immediately after that, sevoflurane administration was stopped, and 100% oxygen was administered until the ewe was awake. After confirming the anesthesia level above the midline laparotomy, lambs were delivered. Amniotic fluid and umbilical cord blood samples were taken at the delivery. After delivery, the ewes were euthanized with intravenous pentobarbital (140 mg/kg; Euthasol, Pharma, Oudewater, Netherlands). The lambs were stimulated and dried with towels on a heating pad and under an infrared lamp. After regular breathing was established, a 22-gauge catheter was inserted in the right jugular vein. The lambs were then transferred for magnetic resonance imaging (MRI). Thereafter, 5 mg boluses of propofol (PropoVet multidose, Ecuphar, Oostkamp, Belgium) were repeated through the jugular vein catheter until lamb pupil reflexes disappeared. The bowel was then removed for obduction, and the lambs were eventually euthanized with intravenous sodium pentobarbital (140 mg/kg).
Following ultrasound confirmation of fetal cardiac activity, *mid-gestation* fetal lambs (13–21 days post-induction) underwent laparotomy with uterus exteriorization and ultrasound assessment under general anesthesia followed by collection of amniotic fluid and umbilical cord blood via hysterotomy. Ewes and fetuses were subsequently euthanized with maternal intravenous administration of pentobarbital. Upon delivery, a complete necropsy was performed, and bowel specimens were collected for detailed macroscopic and histological examination.

**In vivo MRI bowel morphology and motility**
In live *term* neonatal lambs, an abdominopelvic MRI was performed within two hours after delivery using a 3.0 Tesla MRI (Magnetom Prisma, Siemens Healthcare, Erlangen, Germany) with an 18-channel body coil. During the 30–45 min scanning period, lambs were continuously monitored and received intermittent intravenous boluses of 5 mg propofol as needed to minimize motion. Three gradient-echo MRI sequences were acquired: (1) TRUe Fast Imaging with steady-state free precession (TRUFI) sequences to enhance intraluminal fluid contrast of the small bowel, (2) TRUFI cine sequences to capture small bowel motility, and (3) T1-weighted Volumetric Interpolated Breath-hold Examination (VIBE) sequences before and after intravenous gadolinium contrast (2 mL, 0.5mmol/L; Dotarem, Guerbet, Villepinte, France) administration, to precisely visualize the bowel wall and to determine bowel wall thickness. All sequences were acquired in three orthogonal planes (axial, coronal, and sagittal). For the cine imaging, 3-mm-thick volume blocks were acquired every second over a 2-min period, resulting in three slices with 100 images per slice.
Structural parameters included intra-abdominal bowel dilation and intra-abdominal bowel wall thickness, while functional assessment comprised GIQuant motility and gastrointestinal (GI) contrast scores. Structural measurements were performed using Horos (version 3.3.6; Horos project, Brooklyn, NY, USA). For intra-abdominal bowel dilation, the three most dilated bowel segments were identified anterior to the left kidney on TRUFI sequences in each plane, and diameters were measured inner-to-inner wall (mm). The mean of three measurements per plane and across planes was calculated. For intra-abdominal bowel wall thickness, three different bowel segments were selected anterior to the left kidney on axial T1-weighted VIBE images after contrast administration. Bowel wall thickness was measured outer-to-inner wall (mm), and the mean of the three measurements was taken.
GIQuant motility score and GI contrast score were derived from cine sequences using the GIQuant tool (Motilent, London, UK), a well-validated cine-MRI analysis platform.^5-8^ GIQuant motility score quantifies bowel wall motion based on spatiotemporal signal variation, whereas GI contrast score quantifies intraluminal signal retention over time as a surrogate of intestinal transit. Regions of interest (ROI) were manually delineated using a smooth polygon with a minimum area of 1 cm² on a representative frame of the cine sequence on Entrolytics (Motilent, London UK). These ROI were applied to the corresponding cine-derived heatmaps, from which GIQuant motility scores were calculated, while GI contrast scores were derived from the cine sequences. In non-operated littermates, a single ROI was defined in small bowel loops anterior to the left kidney and used for both GIQuant and GI contrast analysis. In gastroschisis-induced lambs, separate ROI were defined: one in exteriorized bowel for GIQuant motility assessment and another in intra-abdominal bowel to assess intestinal transit proximal to the abdominal wall defect (**Figure S1**). Mean GIQuant motility and GI contrast scores were calculated across the three planes.

**Tissue collection and preparation**
Fresh tissues were collected in a standardized manner. Bowel specimens were sampled from the proximal jejunum, jejunoileal junction, and terminal ileum.^9^ The proximal jejunum was taken from the bowel part with arterial supply from the first two branches of the CMA and was used for histology. The jejunoileal junction was localized based on the two middle branches of the CMA and was used for ex vivo bowel contractility testing, ex vivo intestinal permeability testing, and histology. The terminal ileum was identified based on the two distal branches of the CMA and was used for histology.^10-12^ The remaining three bowel segments were individually stored at -80˚C for subsequent omics analyses.
Amniotic fluid, maternal blood, and umbilical cord blood samples collected at delivery, both at *term* and *mid-gestation*, were immediately centrifuged for 10 min at 3,000 rpm using a Centrifuge 5703 (Eppendorf AG, Hamburg, Germany). The supernatant was aliquoted into cryogenic storage tubes (Thermo Fisher Scientific, Waltham, MA, USA) and stored at −80 °C until analysis.
The remaining bowel segments and blood samples were collected and stored prospectively but were not analyzed in the present study.

**Ex vivo bowel contractility testing**
Ex vivo bowel contractility testing was performed within one hour after euthanasia. First, the jejunoileal specimens were rinsed with ice-cold saline, then full-thickness strips (length 10 mm, width 2 mm in *term* lambs) or segments (length 10 mm in *mid-gestational* fetuses) were suspended and stretched (0.2–0.5 gF) along their longitudinal axis, in an organ bath maintained at 37 ˚C and filled with Krebs solution (119.03 mmol/L NaCl, 2.0 mmol/L NaH_2_PO_4_, 15.5 mmol/L NaHCO_3_, 5.9 mmol/L KCl, 2.50 mmol/L CaCl_2_, 1.2 mmol/L MgCl_2_ and 11.5 mmol/L glucose). The organ bath was continuously aerated with a gas mixture of 95% O_2_/5% CO_2_, warmed and humidified through contact with the 37 ˚C Krebs solution. Mechanical responses in the smooth muscle strips were measured using an isometric force transducer/amplifier (Harvard Apparatus Inc., South Natick, MA, USA) and a DI-720 data acquisition system with Windaq software (Dataq Instruments, Akron, OH, USA). After stabilization (1h) at optimal stretch, acetylcholine (ACh) (100 µM) was added until a stable contraction was obtained, followed by a 20-min washout period. This was repeated until a stable response to ACh was obtained, typically two to three times. Next, the strips were challenged with a buffer containing 60 mM potassium chloride (KCl), followed by a 30-min washout. Finally, 100 nM substance P was added to the bath with a 30-min washout period afterwards. Neural responses were elicited using electrical field stimulation (EFS) applied via two parallel platinum rod electrodes using a Grass S88 stimulator (Grass Technologies, West Warwick, RI, USA). Frequency spectra (0.5–32 Hz) were established using pulse trains with a pulse duration of 0.35 ms, a train duration of 10 seconds, and an amplitude of 10 V. Voltage was kept constant by using a Med Lab Stimu-Splitter II (Med Lab, Loveland, CO, USA). A 90-second interval was left after each pulse train. The neural responses during stimulation (ON responses) were calculated as the mean contractile response above baseline during the stimulation period. Post-stimulation responses (OFF responses) were recorded but were not included in the analyses presented in the main manuscript. The baseline tone was defined as the median response during 30 seconds before the stimulation period. Contractile responses were expressed as force normalized to tissue cross-sectional area (gF/mm^2^), with KCl used as a viability control. The pharmacology of the neural responses was assessed by performing EFS under non-adrenergic, non-cholinergic (NANC) conditions (5 µM atropine, 500 μM guanethidine), followed by sequential addition of L-NAME (300 μM) and MRS-2500 (1 μM).
*Data analysis*
Individual basal EFS frequency-response curves were fitted using a four-parameter logistic model^13^ with the lower and upper asymptotes fixed to each lamb’s observed minimum and maximum responses. The stimulation frequency required to achieve 50% of the maximal neural response (half-maximal frequency) and slope factor were estimated by minimizing the sum of squared residuals using Excel Solver (GRG Nonlinear), constrained to a half-maximal frequency of 0.5–32 Hz and a slope factor ≥0. Fits were initialized from two predefined starting values (half-maximal frequency 1 Hz, slope factor 1; and half-maximal frequency 4 Hz, slope factor 2) to assess convergence. A half-maximal frequency was considered non-estimable when a stable solution within the predefined parameter space could not be obtained. Half-maximal frequencies were log₂-transformed for statistical analysis.
The maximal neural response was defined as the contractile response at the highest stimulation frequency (32 Hz). Pharmacological effects were quantified as incremental changes between sequential conditions at 16 Hz: the NANC effect was calculated as the response under NANC conditions minus the basal EFS response; the L-NAME-associated effect as the response under NANC + L-NAME minus the response under NANC conditions; and the MRS-2500-associated effect as the response under NANC + L-NAME + MRS-2500 minus the response under NANC + L-NAME.
Log₂-transformed half-maximal frequencies, maximal neural responses, and pharmacological effects were compared according to gastroschisis type at necropsy, using the Mann–Whitney U for two-group comparisons and the Kruskal–Wallis with Dunn’s post hoc test for three-group comparisons. For graphical representation, group-level four-parameter logistic curves were generated using the mean fitted parameters for each group. Mean observed responses at each stimulation frequency are shown with error bars representing the standard error of the mean (SEM); these graphical representations are descriptive rather than inferential given the small sample size.

**Ex vivo intestinal permeability testing**
Ex vivo intestinal permeability testing was conducted within one hour after euthanasia; due to feasibility, this was only attempted in term lambs. Intestinal segments (10 x 2 mm) were collected from the jejunoileal junction, rinsed in phosphate-buffered saline (PBS; pH 7.4, prepared in-house according to standard protocols), and the seromuscular layer was surgically removed. The mucosa was mounted in a modified Ussing chamber (Mussler Scientific Instruments, Simmerath, Germany) using sliders with a 0.49-cm^2^ aperture (#P2305, Physiologic Instruments, Venice, FL, USA). The luminal and basolateral compartments were filled with Krebs-Ringer bicarbonate buffer supplemented with 10 mmol/L glucose. Solutions were kept at 37°C and continuously gassed with a mixture of 70% oxygen and 30% carbondioxide. Data were acquired using Acquire & Analyze data acquisition software (Acquire & Analyze II, Physiological Instruments, Venice, FL, USA). To assess epithelial integrity, transepithelial electrical resistance (TEER) was measured. Transmucosal potential difference was continuously monitored using Ag/AgCl electrodes. TEER was calculated using Ohm’s law from the voltage deflections induced by bipolar current pulses of 50 μA applied every 60 seconds, each lasting 200 ms under short-circuit conditions. Measurements were collected at 0, 30, 60, 90, and 120 min and reported in Ohm*cm^2^. To assess intestinal permeability, fluorescein isothiocyanate (FITC) permeability testing was conducted. Molecular flux was studied by adding 4 kDa dextran conjugated with FITC (FITC-Dx4, 1 mg/mL, Sigma-Aldrich, Hoeilaart, Belgium) to the luminal compartment after a 40-min stabilization period. Fluorescence was determined in samples from the basolateral side, taken at 0, 30, 60, 90, and 120-min time points. Fluorescence values were converted to picomol/cm^2^ based on a standard curve generated for each experimental run.^14^
*Data analysis*
Intestinal permeability data were analyzed using mixed-effects modeling to account for repeat measurements over time. Fixed effects included time, group (phenotypes), and their interaction, while random effects accounted for subject variability. For graphical representation, the mean was used for individual data points, and error bars represent the SEM, acknowledging this is descriptive and not inferential due to small sample size.

**Histological assessment of bowel morphology**
Bowel specimens were immersed in 4% paraformaldehyde (PFA; Sigma-Aldrich, St. Louis, MO, USA) for 24 hours, washed with PBS for an hour, and transferred to 70% ethanol (VWR International, Leuven, Belgium) for dehydration. The specimens were cut into three pieces: a middle part was sectioned longitudinally, and the two lateral parts were for transverse sections. Following embedment in paraffin, four µm sections were made with an automatic microtome (HM 355S Automatic Microtome, Thermo Fisher Scientific, Waltham, MA, USA). Sections were stained with hematoxylin and eosin (H&E) and periodic acid Schiff (PAS) or kept for immunohistochemistry; immunohistochemistry was not analyzed in the present study. Images were digitized using ZEISS Axioscan 7 slide scanner (Carl ZEISS Microscopy GmbH, Jena, Germany) and ZEISS ZEN 3.4 (blue edition) software with a maximum resolution of ×40. On H&E slides, 20 well-oriented complete villus-crypt units were selected in the longitudinal section.^12^ Villi length and crypt depth – defined as the distance between the mouth of the crypt and the tip of the villus, and that between the mouth of the crypt and the bottom of the crypt – were measured and adjusted according to the average of the 20 measurements.^10^ When more than 20 complete units were available, the longest 20 villi and corresponding crypts were chosen.^12^ In two transverse sections per lamb, 20 measurements were performed for both the inner circular and outer longitudinal layers of the muscularis externa (40 measurements total).^12^ Thickness values were summed to represent total muscular thickness and also reported separately. These measurements on H&E slides were adjusted for birth weight.^11,12^ Additionally, smooth muscle cell quantification was performed on H&E slides using QuPath. Manually annotated regions of the circular and longitudinal smooth muscle layers were defined to include at least 1200 nuclei. Nuclei were detected using watershed-based cell detection,^15^ and cell density was calculated as the number of detected nuclei per annotation area (#nuclei/1000µm^2^).^16,17^ Results were reported separately for each layer.
On PAS sections, cell-counting macros were defined, including 20 villus-crypt units in term lambs (10–15 in *mid-gestation* fetuses) within the mucosal layer of the bowel in the transverse section (**Figure S2**).^11^ Goblet cells were identified by a vibrant pink/magenta color. Cells were first detected, and their features were extracted using QuPath with StarDist plug-in. Then those cells were classified using a trained object classifier in QuPath.^15,18^ The goblet cell number was shown as a percentage of positively stained cells per all epithelial cells counted.^11,12^

**Amniotic fluid biochemical analysis**
Amniotic fluid samples, previously centrifuged and aliquoted, were stored at −80 °C until analysis. Biochemical analyses were performed at the Department of Laboratory Medicine, University Hospitals Leuven (Herestraat 49, 3000 Leuven, Belgium). Samples were thawed and analyzed on a Roche cobas c702 analyzer (Roche Diagnostics, Basel, Switzerland). The following analytes were measured: total protein using the TPUC3 reagent for the low measurement range (40–2,000 mg/L; Roche Diagnostics, 05171954) and the TP2 reagent for the high measurement range (2–120 g/L; Roche Diagnostics, 05171385); amylase (AMYL2 reagent; Roche Diagnostics, 05167027); lipase (LIPC reagent; Roche Diagnostics, 07041918); bile acids (Diazyme Total Bile Acids Assay Kit; Diazyme Laboratories, DZ042A); and L-lactate (LACT2 reagent; Roche Diagnostics, 05171881). For total protein measurement, the low measurement range was used primarily; the high measurement range was applied when values exceeded the upper limit of the low range.

**Additional methodological considerations**
The experimental unit was the individual fetal lamb. Only surviving fetal lambs at assessment were included in statistical analyses; IUFDs were recorded but excluded from analyses. No other exclusion criteria were applied.
Analyses were omitted when technically infeasible (e.g., insufficient fluid sample volume, apparatus availability, or logistical constraints); therefore, the number of observations is reported for each analysis. Exclusions were based solely on predefined survival criteria or technical feasibility and were independent of outcome measures. Blinding varied by outcome measure: for prenatal ultrasound assessments, the sonographer could not be blinded to group allocation due to the visible presence of gastroschisis, whereas assessors for all other outcomes were blinded to group allocation.
Assessment time points were determined a priori based on planned surgical schedules and logistical feasibility; formal randomization of assessment timing was not performed. Fetuses selected for induction were chosen based on anatomical position, which determined surgical accessibility and procedural risk; therefore, random allocation at the fetal unit level was not feasible.

**Supplementary figure legends**

**Figure S1. In vivo MRI assessment of bowel motility.**
TRUFI cine sequences were used to assess bowel motion. ROI (>1 cm^2^) were manually delineated on a representative frame of the cine sequence (A, C). Corresponding GIQuant bowel heatmaps are shown in (B, D), from which GIQuant motility scores were calculated, while GI contrast scores were derived from the cine sequences.
Panels (A, B) show a non-operated lamb, and (C, D) a gastroschisis-induced lamb. In non-operated lambs, a single ROI was used for both GIQuant and GI contrast analysis. In gastroschisis-induced lambs, separate ROI were defined in exteriorized bowel for GIQuant motility assessment and in intra-abdominal bowel for GI contrast analysis to evaluate intestinal transit proximal to the abdominal wall defect.
Abbreviation: GI, gastrointestinal; ROI, regions of interest; TRUFI: true fast imaging with steady-state free precession.

**Figure S2. Goblet cell quantification.**
Upper: Periodic acid-Schiff staining highlighting goblet cells (magenta) in the intestinal mucosa. Lower: Automated goblet-cell detection using QuPath with the StarDist plugin. Cell-counting macros were applied manually across 20 villus-crypt units (10–15 in mid-gestation fetuses). *Scale bar: 200 µm.*

**Figure S3. Additional representative necropsy findings.**
(A) Bowel stenosis at the level of the silicone ring. (B) Macerated fetus after intrauterine fetal death demonstrating bowel morphology suggestive of volvulus (arrowhead). (C) Externally visible bowel perforation with meconium leakage (arrowhead). (D) Bowel atresia identified during dissection (arrowhead).

**Figure S4. Additional results of ex vivo bowel contractility testing.**
Results are shown for term lambs (A) and mid-gestation fetuses (B). Upper panels show fitted four-parameter logistic curves for contractile responses to EFS. Lower panels summarize isometric contractile responses, derived EFS parameters, and incremental pharmacological responses. ***p*<.01 (Mann–Whitney U test).
Abbreviations: cGS, complex gastroschisis; EFS, electrical field stimulation; freq, frequency; L-NAME, N-nitro-L-arginine methyl ester; MRS-2500, (1’R,2’S,4’S,5’S)-4-(2-Iodo-6-methylaminopurin-9-yl)-1-[(phosphato)methyl]-2(phosphato)bicycle[3.1.0]-hexane; NANC, non-adrenergic non-cholinergic; sGS, simple gastroschisis.

**Figure S5. Ex vivo intestinal permeability assessment in term lambs.**
(A) TEER and (B) FITC-Dx4 flux measured in jejunoileal junction segments at 0, 30, 60, 90, and 120 minutes.
Abbreviations: cGS, complex gastroschisis; FITC-Dx4, fluorescein isothiocyanate-dextran 4 kDa; TEER, transepithelial electrical resistance.

**Figure S6. Additional histological assessment of bowel morphology in term lambs.**
Morphometric measurements were performed in proximal, middle, and distal bowel segments. **p*<.05, ***p*<.01, ****p*<.001 (Mann–Whitney U test).
Abbreviations: BW, birth weight; cGS, complex gastroschisis.

**Figure S7. Histological assessment of bowel morphology in mid-gestation fetuses.**
Morphometric measurements were performed in proximal, middle, and distal bowel segments. Birth weight was unavailable for one complex gastroschisis fetus and two non-operated littermates; therefore, these were excluded from analyses requiring birth weight adjustment.
Abbreviations: BW, birth weight; cGS, complex gastroschisis; sGS, simple gastroschisis.

**Figure S8. Prenatal ultrasound measurements in mid-gestation fetuses.**
Prenatal ultrasound measurements: (A) UA-PI, (B) DV-PI, (C) CMA-PI, (D) IABD, (E) IABWT, (F) EABD, (G) EABWT, and (H) EABV. In panel H, complex gastroschisis is further subdivided into perforated and non-perforated cases. Significance indicators are shown only for comparisons between complex and simple gastroschisis when significant. **p*<.05 (Mann–Whitney U test).
Abbreviations: cGS, complex gastroschisis; CMA, cranial mesenteric artery; DV, ductus venosus; EABD, extra-abdominal bowel dilation; EABV, extra-abdominal bowel volume; EABWT, extra-abdominal bowel wall thickness; IABD, intra-abdominal bowel dilation; IABWT, intra-abdominal bowel wall thickness; NP, non-perforated; P, perforated; PI, pulsatile index; sGS, simple gastroschisis; UA, umbilical artery.

**Figure S9. Exploratory logistic regression and ROC analysis of IABD for prediction of complex gastroschisis.**
ROC analysis evaluated the association between IABD and occurrence of complex gastroschisis. The Youden index identified an optimal probability cut-off of 0.657, corresponding to an IABD of 2.77 mm (rounded to 2.8 mm), with a sensitivity of 0.75 and specificity of 1.00.
Abbreviations: GS, gastroschisis; IABD, intra-abdominal bowel dilation; ROC, receiver operating characteristic.

**Figure S10. Amniotic fluid analysis in mid-gestation fetuses.**
Amniotic fluid concentrations of total protein (A), amylase (B), lipase (C), bile acids (D), and L-lactate (E). Significance indicators are shown only for comparisons between complex and simple gastroschisis when significant. **p*<.05 (Mann–Whitney U test).
Abbreviations: cGS, complex gastroschisis; sGS, simple gastroschisis.

**Supplementary video legend**

**Videoclip S1. Surgical induction of gastroschisis in fetal lambs.**

Creation of a standardized abdominal wall defect, silicone ring fixation, and exteriorization of bowel during fetal surgery at gestational day 75. *Video shown at 2× speed.*

**Supplementary tables**

**Table S1. Baseline characteristic and macroscopic findings of gastroschisis-induced fetal lambs.**

| Fetal ID | Induction (GD) | Assessment (GD) | Interval (days) | Birth weight | Diagnosis | Complex features | Other findings |
| --- | --- | --- | --- | --- | --- | --- | --- |
| Term lambs (n = 14 gastroschisis-induced animals) | | | | | | | |
| T8 | 72 | 77 | 5 | N/A | IUFD | n.r.o. | Maceration |
| T12 | 75 | 99 | 24 | 1347 | IUFD | Volvulus | Maceration, thick fibrotic peel, bowel dilation |
| T13 | 76 | 139 | 63 | 2961 | Complex | Stenosis | Thick fibrotic peel, bowel dilation |
| T14 | 76 | 139 | 63 | 2094 | Complex | Stenosis | Thick fibrotic peel, bowel dilation |
| T10 | 73 | 138 | 65 | 4180 | Complex | Stenosis | Thick fibrotic peel, bowel dilation |
| T11 | 76 | 141 | 65 | 3110 | Complex | Stenosis | Thick fibrotic peel, bowel dilation |
| T1 | 73 | 139 | 66 | 2550 | Complex | Stenosis | Thick fibrotic peel, bowel dilation |
| T2 | 73 | 139 | 66 | 500 | IUFD | n.r.o. | Maceration |
| T5 | 76 | 142 | 66 | 4120 | Complex | Atresia, perforation, necrosis | Thick fibrotic peel, bowel dilation |
| T6 | 72 | 138 | 66 | 4600 | Complex | Stenosis | Thick fibrotic peel, bowel dilation |
| T7 | 72 | 138 | 66 | 1480 | IUFD | n.r.o. | Maceration |
| T9 | 73 | 140 | 67 | 1340 | IUFD | n.r.o. | Maceration |
| T4 | 76 | 144 | 68 | 680 | IUFD | n.r.o. | Maceration |
| T3 | 73 | 142 | 69 | 620 | IUFD | n.r.o. | Maceration |
| Mid-gestation fetuses (n = 18 gastroschisis-induced animals) | | | | | | | |
| M5 | 75 | 88 | 13 | 409 | Simple | No | No peel, small omental cyst, bowel dilation |
| M6 | 75 | 88 | 13 | 400 | Simple | No | Thin peel, bowel dilation |
| M18 | 78 | 91 | 13 | 369 | Complex | Stenosis, atresia, perforation | Partially thick fibrotic peel, bowel dilation |
| M7 | 78 | 92 | 14 | 693 | IUFD | n.r.o. | Maceration |
| M8 | 78 | 92 | 14 | 360 | Simple | No | Thin peel, huge omental cyst |
| M13 | 74 | 88 | 14 | 517 | Simple | No | Thin partial peel |
| M14 | 74 | 88 | 14 | 471 | Simple | No | Partially thick fibrotic peel, small omental cyst |
| M17 | 79 | 94 | 15 | 621 | Simple | No | Thin peel, huge omental cyst |
| M3 | 73 | 89 | 16 | 425 | Simple | No | Partially thick fibrotic peel |
| M4 | 73 | 89 | 16 | 510 | Simple | No | Thin peel |
| M16 | 73 | 90 | 17 | 623 | Simple | No | Thin partial peel |
| M15 | 73 | 91 | 18 | 536 | Complex | Atresia, perforation | Thin peel |
| M2 | 76 | 95 | 19 | 610 | Complex | Atresia | Thick fibrotic peel, bowel dilation |
| M9 | 76 | 96 | 20 | 906 | Complex | Atresia | Thin peel, huge omental cyst, bowel dilation |
| M10 | 76 | 96 | 20 | 935 | IUFD | n.r.o. | Maceration, thin peel |
| M11 | 77 | 97 | 20 | 805 | Simple | No | Thin peel |
| M12 | 77 | 97 | 20 | 894 | Complex | Perforation | Thin peel, huge omental cyst, bowel dilation |
| M1 | 73 | 94 | 21 | N/A | Complex | Atresia | Thick fibrotic peel, bowel dilation |

Fetuses were shown in order of the induction-to-assessment intervals.

Abbreviations: GD, gestational day; IUFD, intrauterine fetal death; M, mid-gestation; N/A, not available; n.r.o., not ruled out; T, term.

**Table S2. Availability of secondary outcome measurements across groups.**

(A) *Term* lambs

|  | Complex GS (n=7) | Normal (n=7) |
| --- | --- | --- |
| In vivo MRI | 5^*^ | 6^*^ |
| Ex vivo bowel contractility | 4^*^ | 6^*^ |
| Ex vivo intestinal permeability | 5^*^ | 6^*^ |
| Histology | 7 (6^‡^ for terminal ileum) | 7 |

(B) *Mid-gestation* fetuses

|  | Complex GS (n=6) | Simple GS (n=10) | Normal (n=10) |
| --- | --- | --- | --- |
| In vivo ultrasound | 4^†^ | 5^†^ | 4^†^ (3^†^ for CMA and IABD) |
| Ex vivo bowel contractility | 1^*^ | 4^*^ | 3^*^ |
| Histology | 5^‡^ (4^‡^ for proximal jejunum and jejunoileal junction) | 10 | 10 |
| Amniotic fluid | 4^‡^ | 5^‡^ | 3^‡^ |

IUFD cases were excluded.

*: Hardware was not available

†: Sample was not taken at the initial phase of the experiment

‡: Sample volume was not sufficient

Abbreviations: GS, gastroschisis; CMA, cranial mesenteric artery; IABD, intra-abdominal bowel dilation; IUFD, intra-uterine fetal death.

**Table S3. Inter- and intra-rater reliability of ultrasound bowel measurements.**

IABD: n=12 images (from 12 fetuses)

| Correlation | Meaning | Estimate | Standard error | 95% confidence interval | P value |
| --- | --- | --- | --- | --- | --- |
| ρ_12_ | Intra-rater (rater 1) | 0.9414 | 0.0343 | [0.9139;0.9603] | <.001 |
| ρ_13_ | Inter-rater (measurement 1) | 0.7045 | 0.1519 | [0.4274;0.8606] | <.001 |
| ρ_23_ | Inter-rater (measurement 2) | 0.7531 | 0.1305 | [0.5308;0.8784] | <.001 |
| ρ | Inter-rater (overall) | 0.7288 | 0.1370 | [0.4885;0.8663] | <.001 |

IABWT: n=12 images (from 12 fetuses)

| Correlation | Meaning | Estimate | Standard error | 95% confidence interval | P value |
| --- | --- | --- | --- | --- | --- |
| ρ_12_ | Intra-rater (rater 1) | 0.9408 | 0.0346 | [0.9129;0.9600] | <.001 |
| ρ_13_ | Inter-rater (measurement 1) | 0.7917 | 0.0112 | [0.7778;0.8048] | <.001 |
| ρ_23_ | Inter-rater (measurement 2) | 0.7175 | 0.1463 | [0.4545;0.8653] | <.001 |
| ρ | Inter-rater (overall) | 0.7546 | 0.1254 | [0.5435;0.8759] | <.001 |

EABD: n=12 images (from 12 fetuses)

| Correlation | Meaning | Estimate | Standard error | 95% confidence interval | P value |
| --- | --- | --- | --- | --- | --- |
| ρ_12_ | Intra-rater (rater 1) | 0.9650 | 0.0243 | [0.9500;0.9755] | <.001 |
| ρ_13_ | Inter-rater (measurement 1) | 0.9872 | 0.0090 | [0.9840;0.9897] | <.001 |
| ρ_23_ | Inter-rater (measurement 2) | 0.9681 | 0.0222 | [0.9552;0.9827] | <.001 |
| ρ | Inter-rater (overall) | 0.9776 | 0.0140 | [0.9711;0.9827] | <0.001 |

EABWT: n=12 images (from 12 fetuses)

| Correlation | Meaning | Estimate | Standard error | 95% confidence interval | P value |
| --- | --- | --- | --- | --- | --- |
| ρ_12_ | Intra-rater (rater 1) | 0.9089 | 0.0615 | [0.8432;0.9478] | <.001 |
| ρ_13_ | Inter-rater (measurement 1) | 0.9186 | 0.0552 | [0.8633;0.9521] | <.001 |
| ρ_23_ | Inter-rater (measurement 2) | 0.9011 | 0.0665 | [0.8526;0.9455] | <.001 |
| ρ | Inter-rater (overall) | 0.9099 | 0.0553 | [0.8526;0.9455] | <.001 |

Reliability was summarized using correlation estimates (ρ) with 95% confidence intervals calculated by Fisher’s z transform (SAS 9.4; SAS Institute, Cary, NC, USA).

Abbreviations: E/IABD: extra-/intra-abdominal bowel dilation, E/IABWT: extra-/intra-abdominal bowel wall thickness.
