## Supplementary figures and images for "A Reproducible Fetal Lamb Model of Complex Gastroschisis with Temporal Characterization of Bowel Changes"

### FigS1

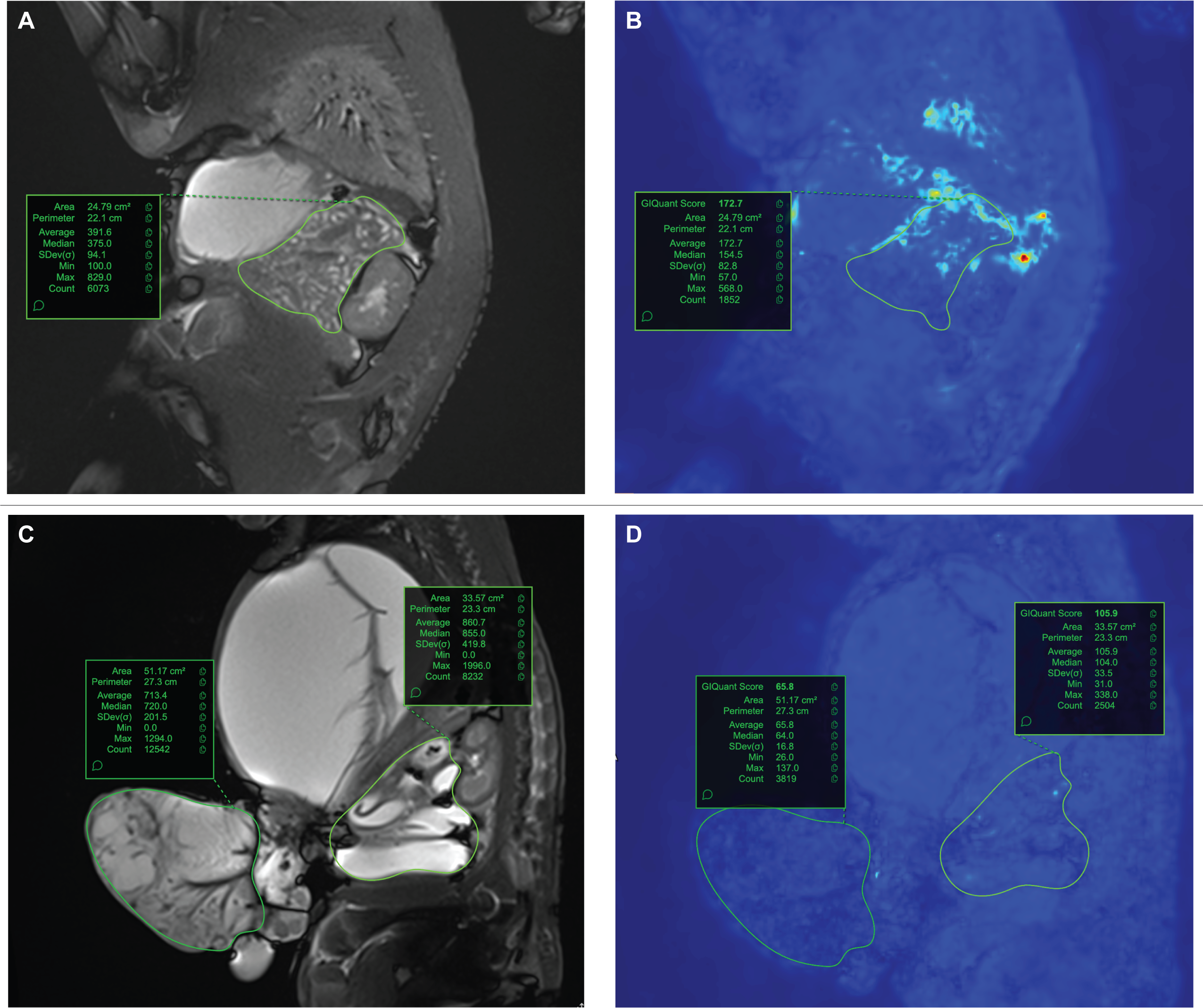

### FigS2

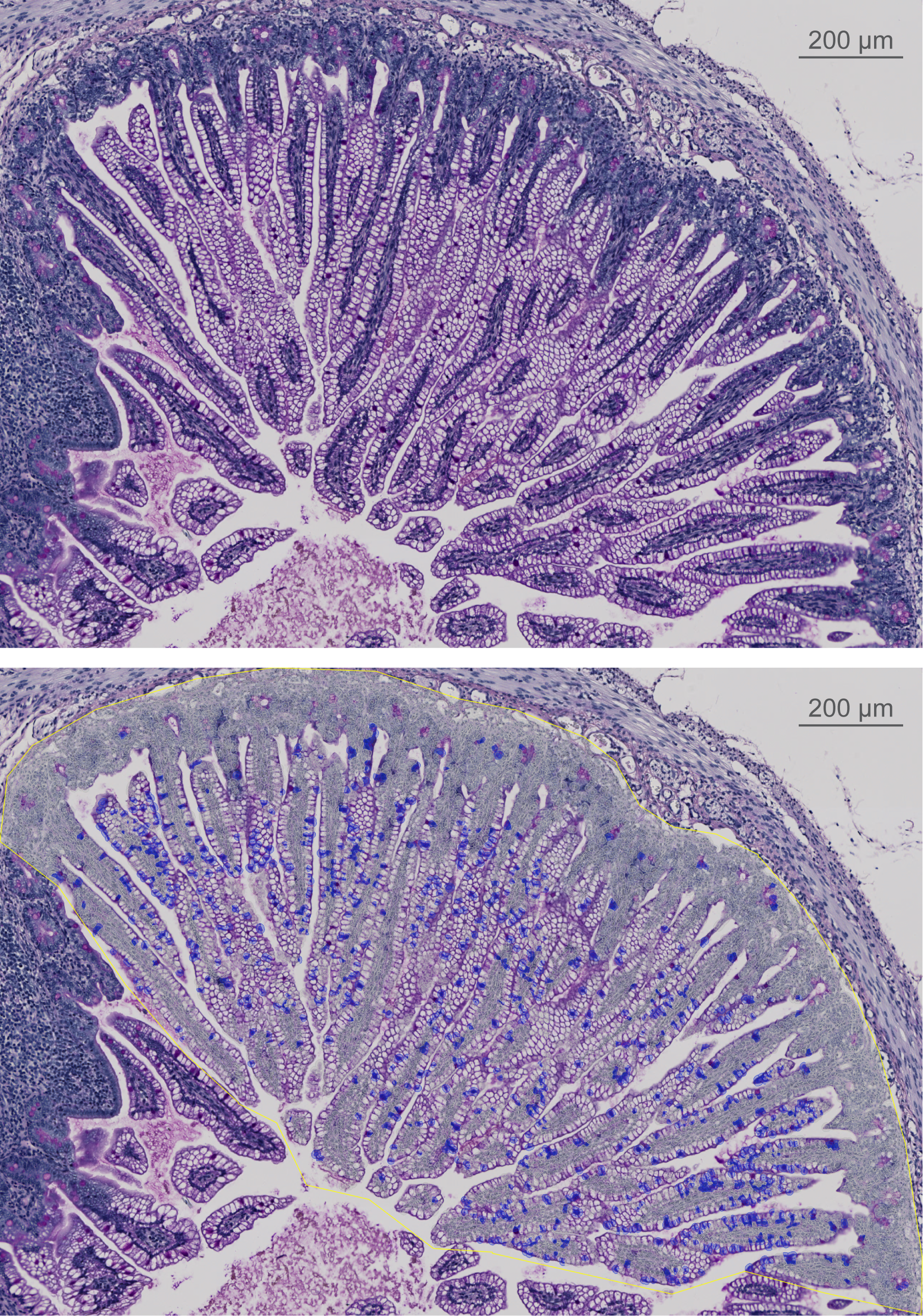

### FigS3

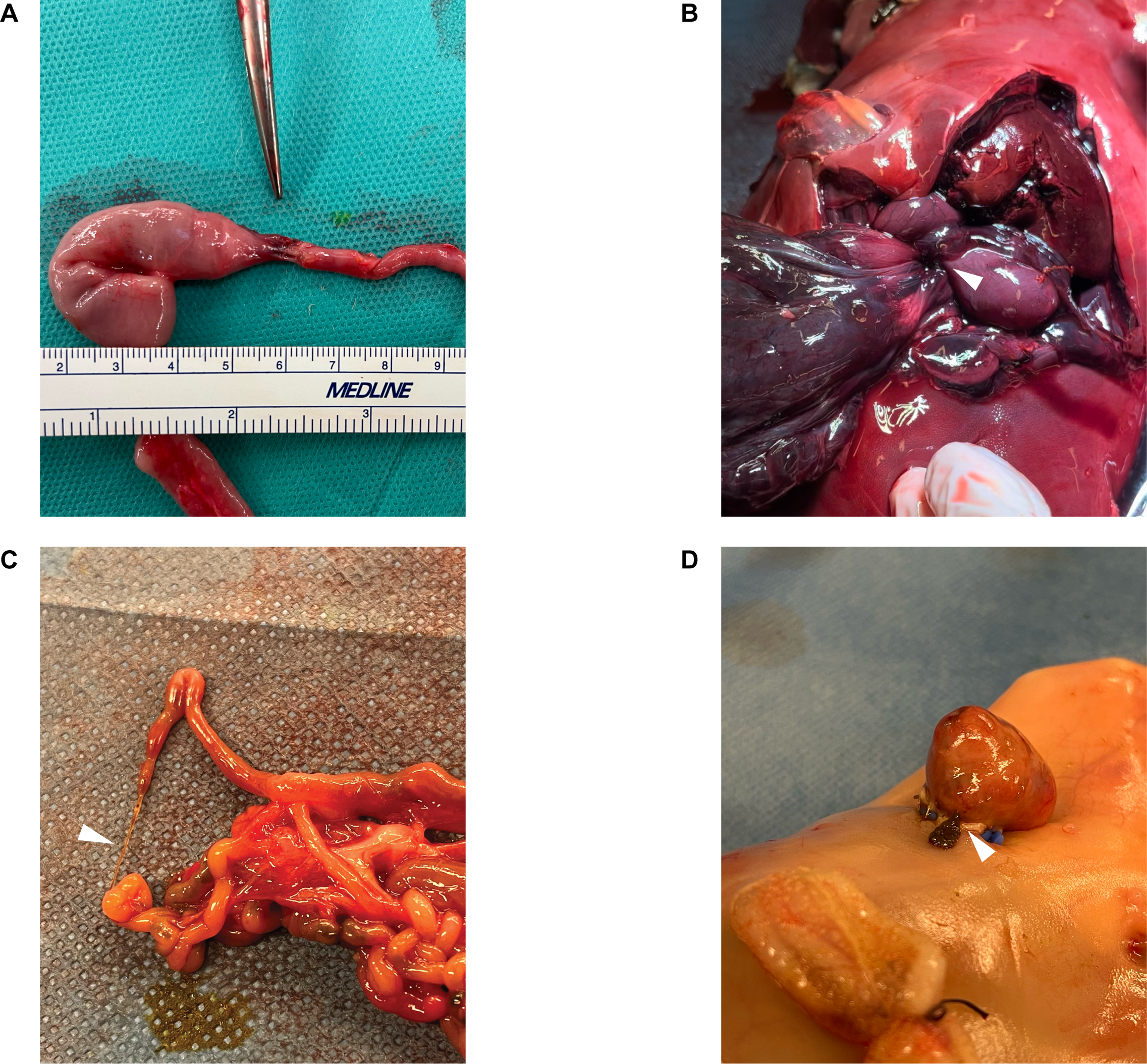

### FigS4

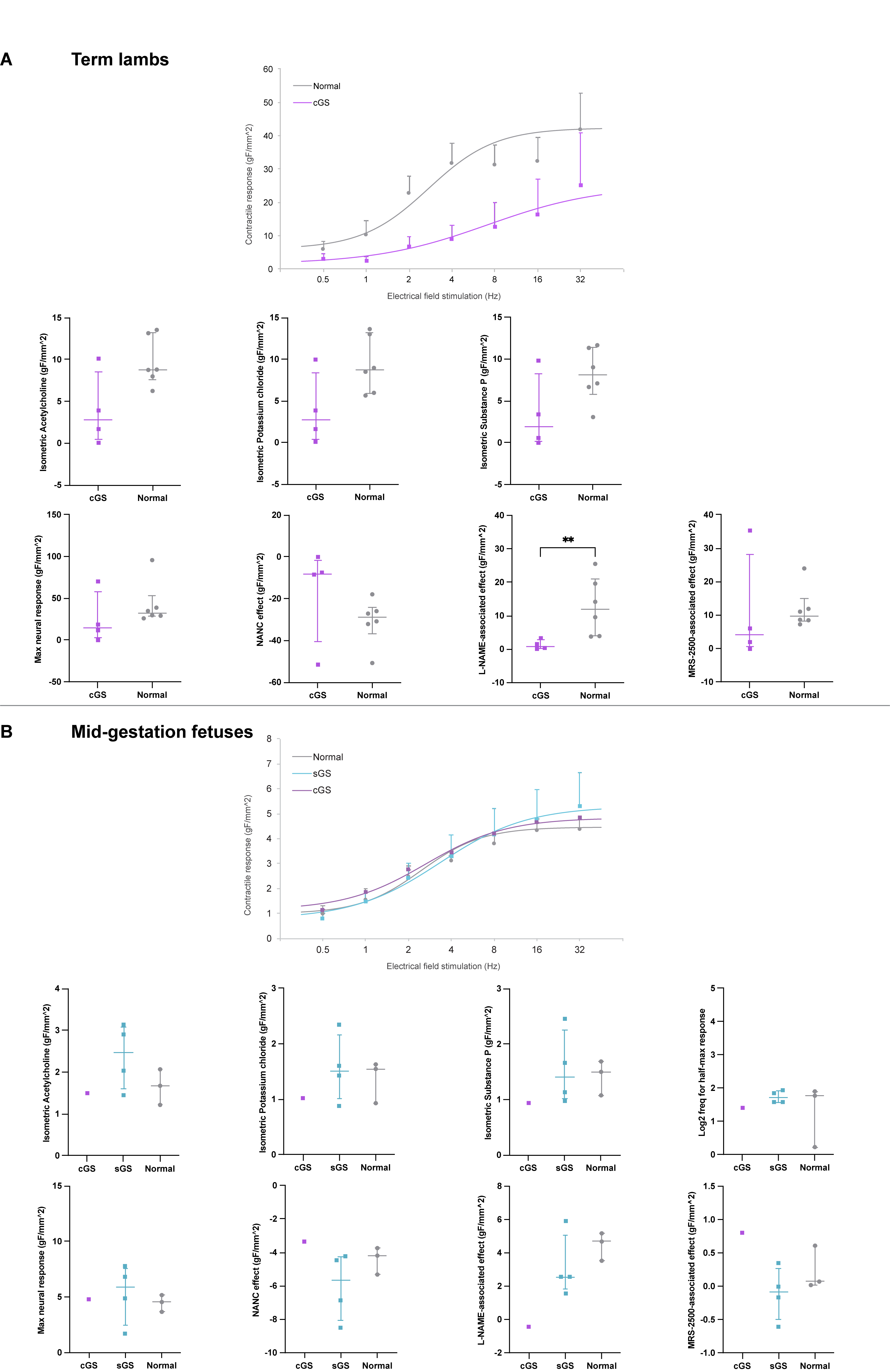

### FigS5

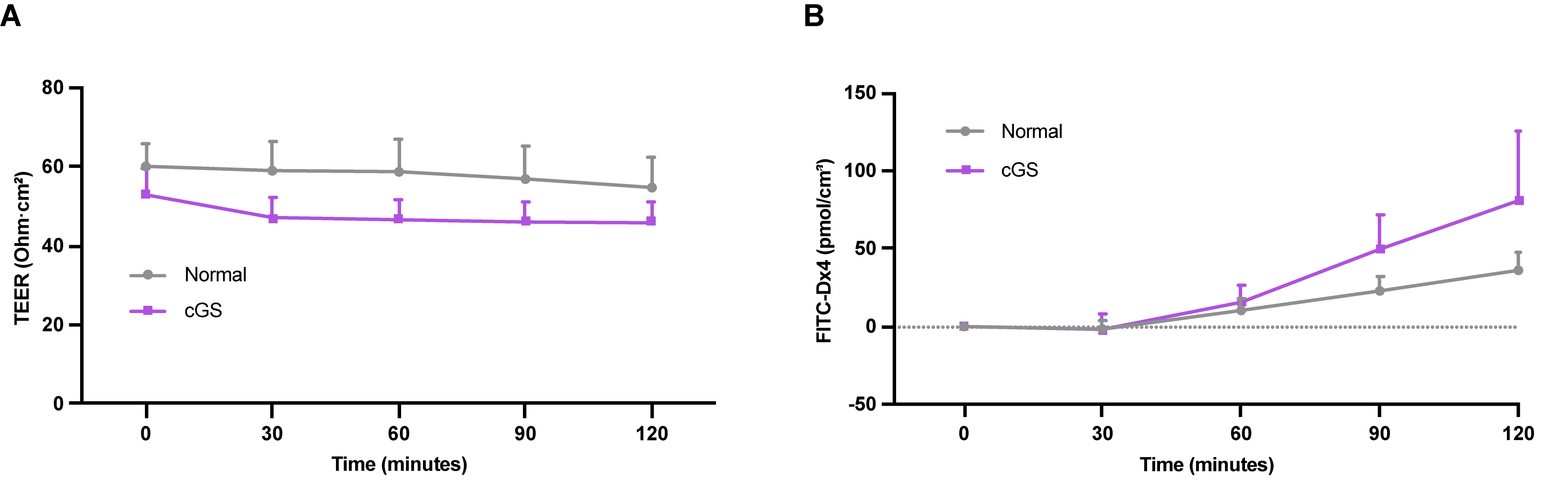

### FigS6

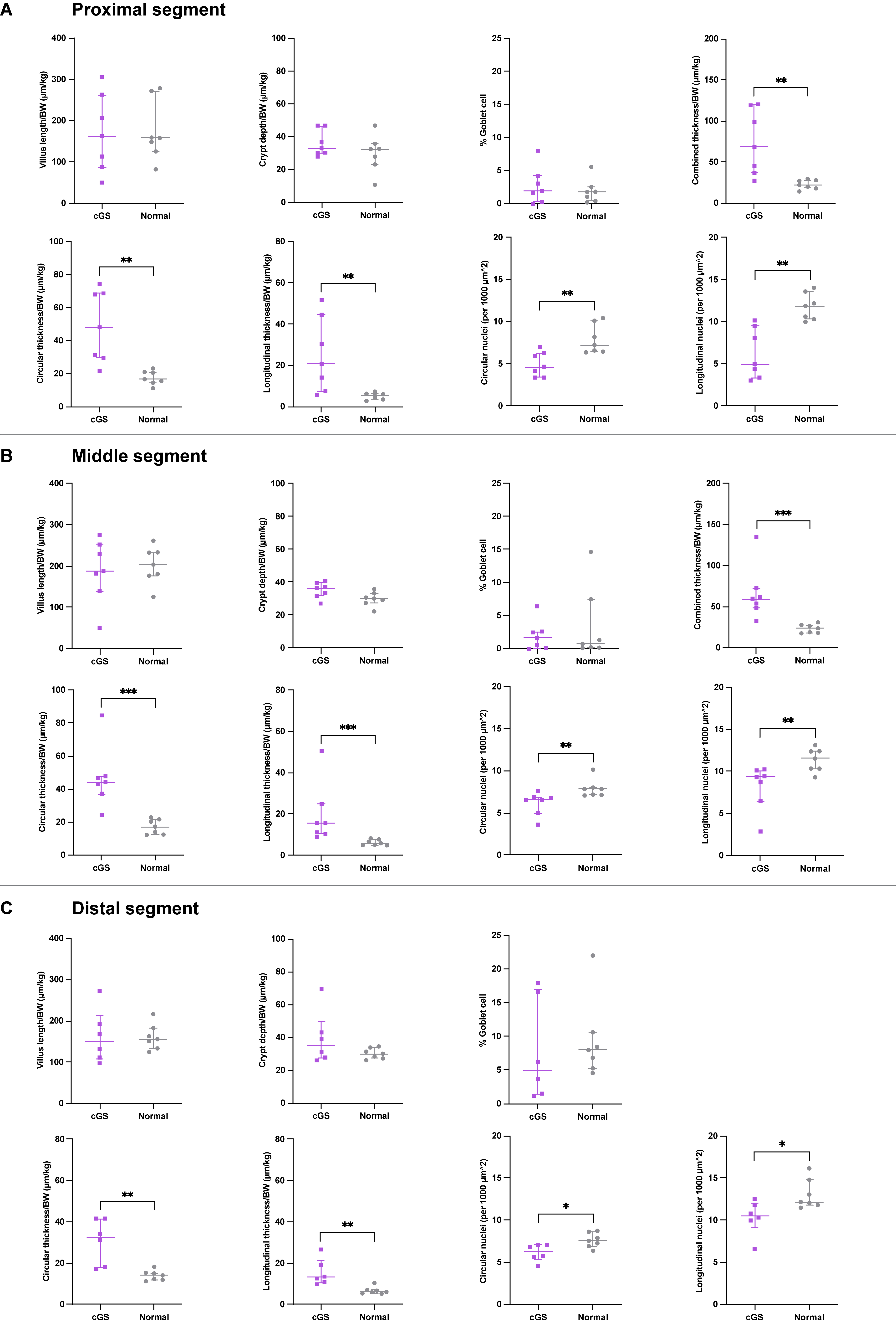

### FigS7

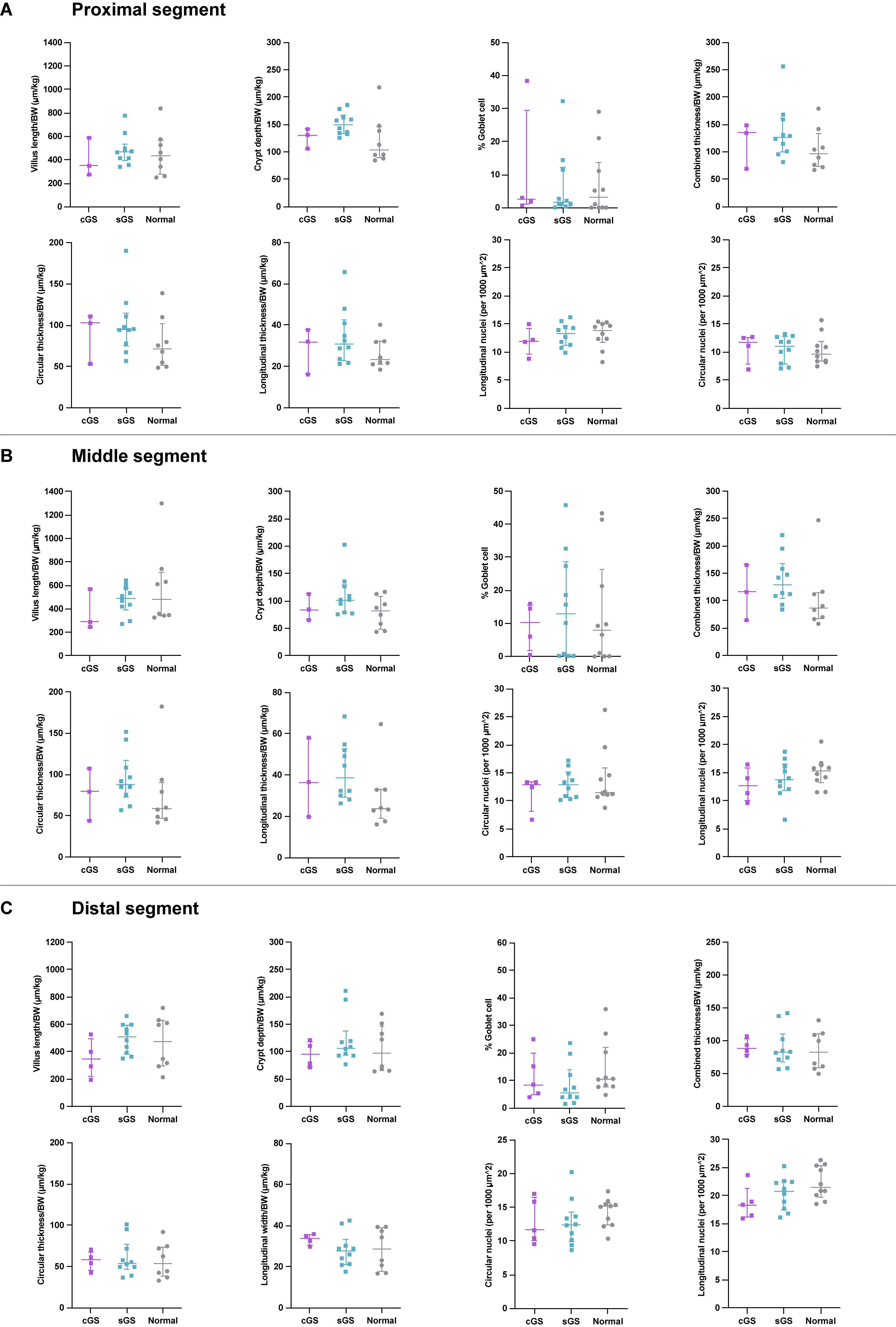

### FigS8

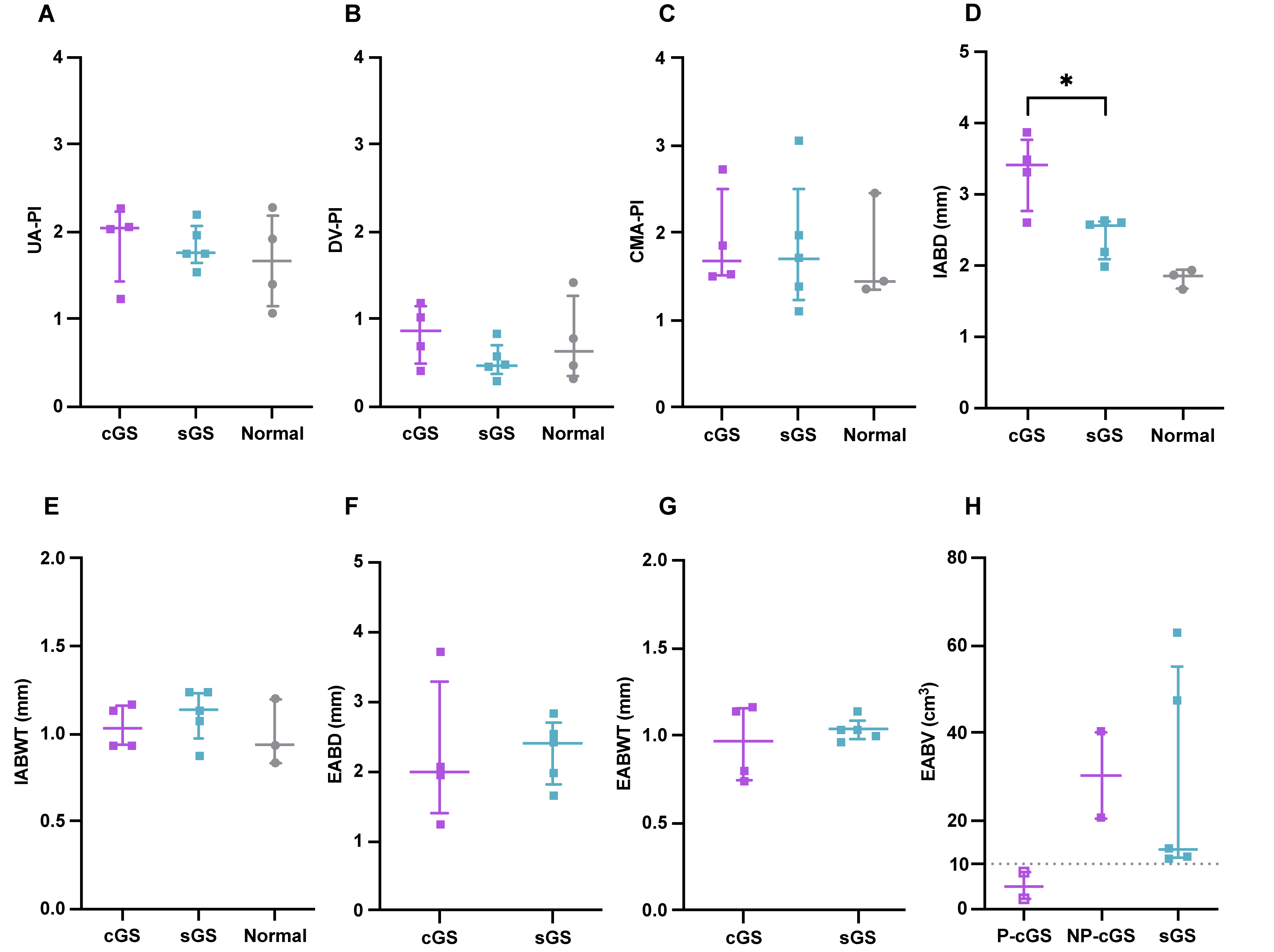

### FigS9

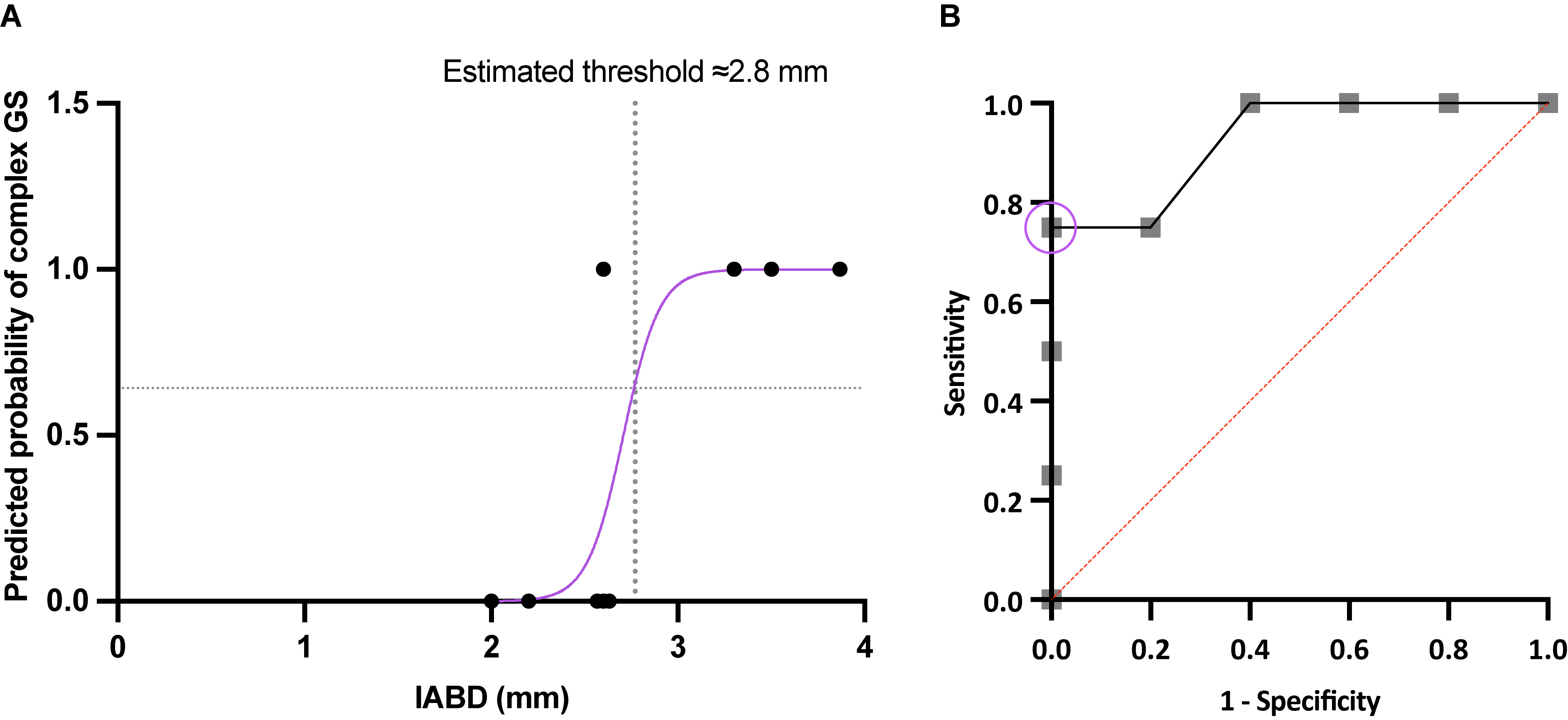

### FigS10

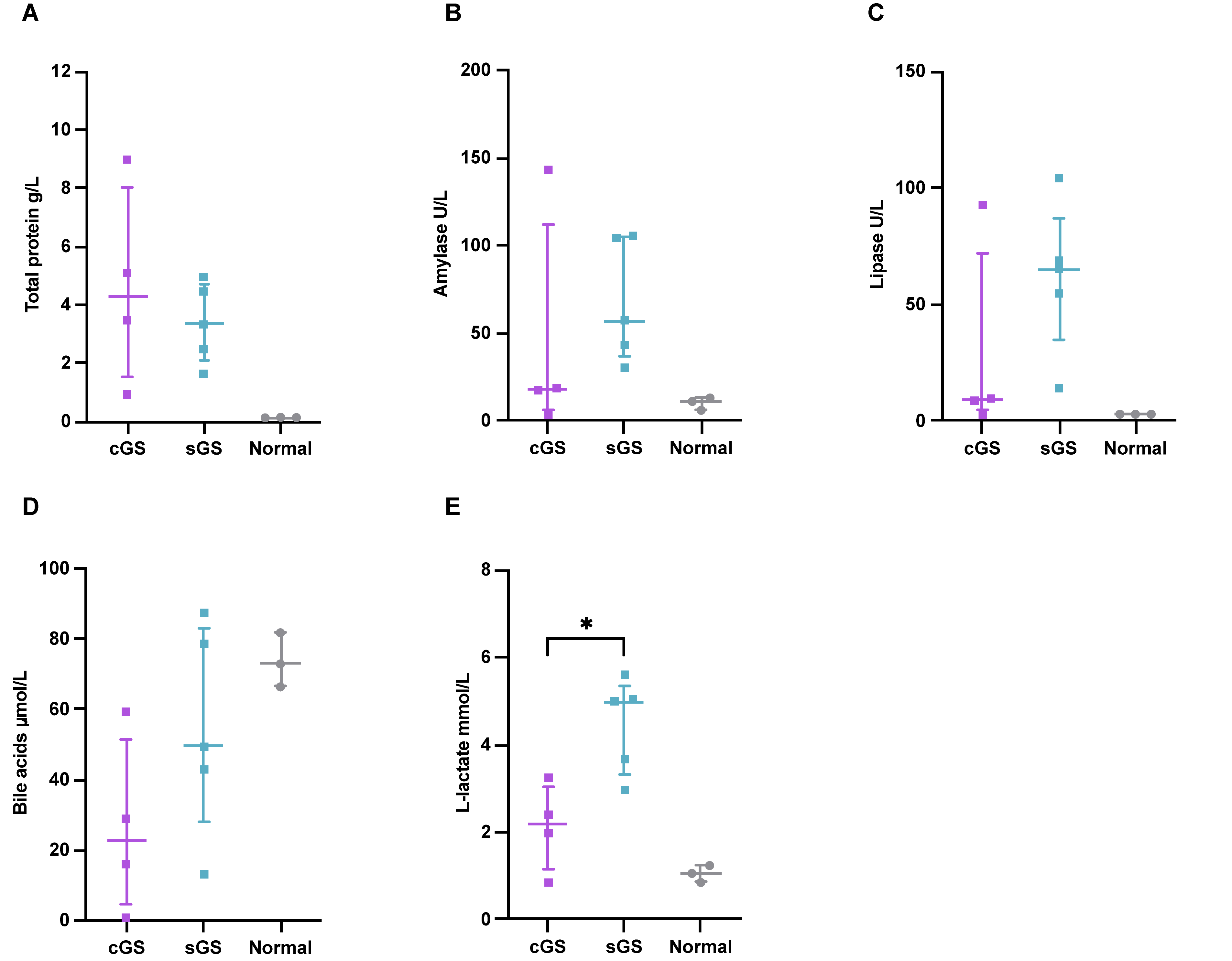
